# Combinations of RAS pathway inhibitors with targeted agents are active in spheroids of patient-derived cells with oncogenic KRAS variants from multiple cancer types

**DOI:** 10.1101/2024.11.17.623998

**Authors:** Zahra Davoudi, Thomas S. Dexheimer, Nathan P. Coussens, Thomas Silvers, Joel Morris, Naoko Takebe, James H. Doroshow, Beverly A. Teicher

**Affiliations:** Molecular Pharmacology Laboratory, Applied and Developmental Research Directorate, Frederick National Laboratory for Cancer Research, Frederick, Maryland, 21702; Division of Cancer Treatment and Diagnosis, National Cancer Institute, National Institutes of Health, Bethesda, Maryland, 20892

## Abstract

The KRAS gene is among the most frequently altered genes in cancer and the KRAS protein was long deemed undruggable. Recent strategies to target oncogenic KRAS have included both direct inhibition of the KRAS protein and indirect inhibition of its activity by targeting upstream and downstream signaling pathway mediators. A high-throughput screen of multi-cell type tumor spheroids was designed to identify active combinations of targeted small molecules and KRAS pathway inhibitors. Inhibitors of the non-receptor protein tyrosine phosphatase SHP2 and the guanine nucleotide exchange factor SOS1 were tested to evaluate indirect upstream pathway inhibition, while sotorasib directly inhibited the KRAS G12C variant. As single agents, sotorasib and the SHP2 inhibitor batoprotafib (TNO155) exhibited selectivity towards spheroids with KRAS G12C, whereas the SOS1 inhibitor BI-3406 showed varying activity across KRAS variants. Vertical inhibition of the RAS/MEK/ERK pathway by targeting SHP2 or SOS1 and the downstream kinases MEK (trametinib) or ERK (temuterkib) was highly effective. Inhibition of upstream tyrosine receptor kinases with nintedanib in combination with batoprotafib or BI-3406 was also effective, and in combination with sotorasib, demonstrated synergy in spheroids harboring KRAS G12C. Dual inhibition of the RAS/MEK/ERK and PI3K/AKT/mTOR pathways with batoprotafib or sotorasib with either the mTORC1/2 inhibitor sapanisertib or the AKT inhibitor ipatasertib demonstrated combination activity, primarily in spheroids harboring KRAS G12C. Combination of the BCL-2 inhibitor venetoclax with sotorasib, batoprotafib or BI-3406 resulted in additive and synergistic cytotoxicity. Lastly, concurrent inhibition of the KRAS pathway with sotorasib and batoprotafib demonstrated combination activity in spheroids containing KRAS G12C.

**SIGNIFICANCE:** KRAS variants are oncogenic drivers for a range of human cancers. Multiple combinations of small molecule agents that target RAS signaling were screened and reduced the viability of multi-cell type spheroid models for a variety of human solid tumors. Combinations warranting further testing were identified.

## INTRODUCTION

RAS (rat sarcoma virus) proteins are a family of small guanosine triphosphate hydrolases (GTPases). Among them, KRAS (Kirsten rat sarcoma viral oncogene homolog) is a frequently altered oncogene in human cancers and is implicated in cancer types with high mortality rates, including pancreatic ductal adenocarcinoma (PDAC), non-small cell lung cancer (NSCLC), and colorectal cancer (CRC) (1-4). As a GTPase, KRAS regulates signal transduction by functioning as a binary switch between its active GTP-bound and inactive guanosine diphosphate (GDP)-bound structures. KRAS signaling is mediated by guanine nucleotide exchange factors (GEFs) and GTPase-activating proteins (GAPs) (3). Gain-of-function variants of the residues G12, G13, and Q61 enhance GTP binding due to an increased nucleotide exchange rate and/or impaired GAP interactions and promote oncogenesis (5). KRAS mediates upstream signals from growth factor receptors such as the epidermal growth factor receptor (EGFR) to downstream signaling pathways, including the mitogen-activated protein kinase (MAPK) signal pathway, the phosphatidylinositol 3-kinase (PI3K)/AKT/mTOR pathway, and the RalGEF/Ral pathway (6-8).

For decades, RAS proteins were considered “undruggable” and RAS has been a challenging drug target due to its small size and apparent absence of allosteric binding pockets in early crystal structures. Additionally, RAS has picomolar affinities for both GDP and GTP which poses a challenge for the development of competitive inhibitors because intracellular concentrations of GTP and GDP are in the micromolar range (9,10). An ability to target the KRAS variant G12C followed the discovery of a novel pocket near the switch regions and proximal to the abnormal cysteine residue. This discovery enabled the development of selective inhibitors that covalently bind the reactive cysteine residue to stabilize the inactive GDP-bound structure and impair interactions with Raf (11-17). Additionally, analogs of GDP were reported that covalently bind and selectively inhibit KRAS G12C (18,19).

The discovery of a hydrophobic switch-II groove on the surface of KRAS G12C (20) enabled the development of the lead compound MRTX1257 and clinical candidates MRTX849 (21) and AMG 510 (22). Sotorasib (AMG 510) was the first KRAS G12C inhibitor to enter a clinical trial (23) and it demonstrated activity in solid tumors with the KRAS G12C variant from heavily pretreated patients (CodeBreaK 100, clinicaltrials.gov; NCT03600883) (24). Long-term clinical studies demonstrated the safety and durable efficacy of sotorasib in patients with pretreated advanced NSCLC harboring KRAS G12C (25) and sotorasib significantly increased progression-free survival compared with the standard of care, docetaxel (clinicaltrials.gov; NCT0430780) (26). In 2021, the U.S. Food and Drug Administration (FDA) granted accelerated approval to sotorasib for the treatment of locally advanced or metastatic NSCLC with KRAS G12C in adult patients (27). Sotorasib, both alone and in combination with other therapeutics, is continuing clinical trial (clinicaltrials.gov; NCT03600883, NCT05180422, NCT04303780, NCT05638295, NCT04625647, NCT05398094, and NCT05118854) (6,28).

As an alternative to direct inhibition of KRAS, several agents have advanced to clinical trials that inhibit factors upstream of RAS and thus might not be affected by RAS status. The GEF SOS1 (Son-of-Sevenless) is required for RAS activation and the Cdc25 domain of SOS1 binds GDP-bound RAS to catalyze the exchange of GDP for GTP (29). SOS1 also interacts with GTP-bound RAS at an allosteric site in the REM (Ras exchanger motif) domain to stabilize the SOS1 active site and augment its GEF function to promote increased catalytic activity (30,31). Structural and functional studies of SOS1 have enabled the development of multiple SOS1 inhibitors including BAY-293, which binds the hydrophobic pocket of the Cdc25 domain and disrupts the interaction between RAS and SOS1 (9,32). The SOS1 inhibitor BI-3406 also binds the Cdc25 hydrophobic pocket and demonstrated improved potency and selectivity compared to previous compounds (33,34). Exposure of cell lines expressing KRAS variants to BI-3406 led to a rapid reduction in GTP-bound KRAS and pERK levels, correlating with growth inhibition (9,34). An analog of BI-3406, BI 1701963, is in clinical trials in patients with advanced solid tumors harboring KRAS variants alone and in combination with agents including irinotecan, the KRAS G12C inhibitor adagrasib, sotorasib or BI 1823911, and the MEK inhibitor trametinib or BI 3011441 (clinicaltrials.gov; NCT04111458, NCT04975256, NCT04835714, NCT04627142, NCT04973163, NCT04185883).

The initiation, propagation, and termination of biological signaling cascades are regulated by protein tyrosine kinases (PTKs) and protein tyrosine phosphatases (PTPs). SHP2 (Src homology-2 domain-containing protein tyrosine phosphatase) is a non-receptor-PTP that is recruited to activated receptor tyrosine kinases (RTKs) and regulates signaling. Activation of the RAS/MEK/ERK signaling pathway by SHP2 involves the dephosphorylation of several negative regulators of RAS, including p120-RasGAP and SPRY/Sprouty to increase GTP-bound RAS (35). Additionally, the SHP2-mediated dephosphorylation of RAS at Y32 enables binding to Raf (36). Inhibitors of the SHP2 PTP catalytic domain have been developed and several are in clinical trials; however, the development of such inhibitors has been challenging due to limited potency, selectivity, permeability, and bioavailability (37). Novartis discovered the first allosteric SHP2 inhibitor SHP836 from a high-throughput screen, which bound in a central tunnel at the interface of the PTP catalytic domain, the N-terminal SH2 domain, and the C-terminal SH2 domain to stabilize an auto-inhibited conformation of SHP2. Medicinal chemistry optimization of SHP836 to SHP099 resulted in a >70-fold increase in potency with improved selectivity and oral bioavailability (38,39). Ultimately this work enabled the discovery of the first-in-class SHP2 inhibitor batoprotafib (TNO155), which is in clinical trials alone and in combinations including KRAS G12C inhibitors such as sotorasib, adagrasib, or JDQ443 (clinicaltrials.gov; NCT03114319, NCT04000529, NCT04185883, NCT04330664, NCT04292119, NCT04294160, NCT04699188, NCT05490030, NCT05541159) (40).

Based on the critical role of the RAS pathway in cellular function and the extensive feedback reactivation in cancers with KRAS variants, a comprehensive blockade of signal transduction is essential for anticancer activity (14,41). Consequently, this study evaluated the KRAS G12C inhibitors sotorasib and MRTX1257 (*data are not shown but are available from PubChem*) as well as indirect KRAS inhibitors BI-3406, BAY-293 (*data are not shown but are available from PubChem*), and batoprotafib alone and in combinations with sixteen FDA-approved and investigational anticancer agents. The single agents and combinations were tested in multi-cell type tumor spheroids, which served as models for human solid tumors and contained malignant cells, endothelial cells, and mesenchymal stem cells (42-44). The spheroids were grown from nineteen malignant cell lines, including patient-derived cell lines from the NCI Patient-Derived Models Repository and an established NSCLC cell line HOP-62. The malignant cell lines were derived from cancers characterized by a high frequency of KRAS alterations, including CRC, NSCLC, and pancreatic cancer.

## MATERIALS AND METHODS

### Compounds

The drugs and investigational agents: BI-3406 (NSC825286), BAY-293 (NSC824723), batoprotafib (TNO155; NSC825523), sotorasib (AMG 510; NSC818433), MRTX1257 (NSC819558), elimusertib (BAY 1895344; NSC800525), molibresib (GSK525762; NSC774829), temuterkib (LY3214996; NSC803410), venetoclax (NSC766270), alisertib (NSC759677), olaparib (NSC753686), talazoparib (NSC767125), erdafitinib (NSC781556), ipatasertib (NSC767898), cabozantinib (NSC761068), nintedanib (NSC756659), abemaciclib (NSC768073), docetaxel (NSC628503), trametinib (NSC758246), and sapanisertib (NSC764658) were obtained from the National Cancer Institute (NCI) Developmental Therapeutics Program Chemical Repository (45). The FDA-approved anticancer drug set is available from the [Developmental Therapeutics Program](https://dtp.cancer.gov/organization/dscb/obtaining/available_plates.htm). The drugs and investigational agents used in this study were demonstrated to be >95% pure by proton nuclear magnetic resonance and liquid chromatography/mass spectrometry. The stock solutions were prepared in dimethyl sulfoxide (DMSO, Sigma-Aldrich, St. Louis, MO, cat. D2650) at 800-fold the tested concentration and stored at -70 °C prior to their use. All drugs and investigational agents were tested over a range starting from a high concentration at or near the clinical C_max_ and decreasing in half-log increments. If the clinical C_max_ for an agent had not been determined, the highest concentration tested was 10 µM (**Table S1**).

### Cell Lines

The patient-derived cancer (PDC) cell lines: 186277-243-T-J2-PDC, 253994-281-T-J1-PDC, 254851-301-R-J1-PDC, 276233-004-R-J1-PDC, 292921-168-R-J2-PDC, 323965-272-R-J2-PDC, 327498-153-R-J2-PDC, 349418-098-R-PDC, 521955-158-R2-J5-PDC, 521955-158-R6-J3-PDC, 519858-162-T-J1-PDC, 885724-159-R-J1-PDC, 931267-113-T-J1-PDC, 941728-121-R-J1-PDC, CN0375-F725-PDC, K00052-001-T-J1-PDC, K24384-001-R-PDC, and LG0567-F671-PDC (**Table S2**) were obtained from the NCI Patient-Derived Models Repository (NCI PDMR). The established cell line HOP-62 was developed by Michael Alley and Michael Boyd from the National Cancer Institute in collaboration with Johns Hopkins University School of Medicine (RRID: CVCL_1285) (**Table S2**) (46). Pooled donor human umbilical vein endothelial cells (HUVEC, Lonza, cat. CC-2519) and human mesenchymal stem cells (hMSC, Lonza, cat. PT-2501) were purchased from Lonza (Walkersville, MD).

### Cell Culture

All cells were maintained in an incubator at 37 °C and 5% CO_2_ with 95% humidity. The PDC lines were cultured according to standard operating procedures established by the NCI PDMR (https://pdmr.cancer.gov/sops/default.htm). Briefly, all PDCs were thawed and cultured in Matrigel-coated flasks prepared with a working solution of 1X Ham’s F-12 nutrient mix, without supplementation (Invitrogen, Waltham, MA, cat. 11765054), 100 U/mL penicillin-streptomycin (Invitrogen, cat. 15140122), and 2.5% Matrigel (Corning Inc., Corning, NY, cat. 354248) for the first three passages. All PDCs were cultured in complete DMEM/F-12 media containing advanced DMEM/F-12 (Invitrogen, cat. 12634028), 4.9% defined fetal bovine serum, heat inactivated (HyClone Laboratories Inc., Logan, UT, cat. SH30070.03HI), 389 ng/mL hydrocortisone (Sigma-Aldrich, cat. H4001), 9.7 ng/mL human EGF recombinant protein (Invitrogen, cat. PHG0313), 23.4 µg/mL adenine (Sigma-Aldrich, cat. A2786), 97.3 U/mL penicillin-streptomycin (Invitrogen, cat. 15140122), 1.9 mM L-glutamine (Invitrogen, cat. 25030081), and 9.7 µM Y-27632 dihydrochloride (Tocris Bioscience, Bristol, United Kingdom, cat. 1254). The PDCs were cultured in complete DMEM/F12 media without 9.7 µM Y-27632 dihydrochloride for at least two passages prior to the screen. The established cell line HOP-62 was cultured in RPMI 1640 medium, HEPES (Invitrogen, cat. 22400105) with 10% defined fetal bovine serum (HyClone Laboratories Inc., cat. SH30070.03). The pooled donor HUVEC were cultured in endothelial cell growth medium 2 (PromoCell, Heidelberg, Germany, cat. C-22011) and the hMSC were cultured in mesenchymal stem cell growth medium 2 (PromoCell, cat. C-28009). For all experiments, HUVEC and hMSC were used at passages ≤5, while the malignant cell lines were used at passages ≤15. Samples of all cell lines were collected at regular intervals throughout the screening process for short tandem repeat (STR) profiling and mycoplasma testing by Labcorp (Laboratory Corporation of America Holdings, Burlington, NC) to confirm their authenticity and integrity.

### High-throughput Drug Combination Screening

Prior to their inoculation into microplates, malignant cells, HUVEC, and hMSC were removed from flasks using TrypLE express (Invitrogen, cat. 12605036) and harvested by centrifugation for 5 min at 233 × g. Following removal of the supernatant, the cells were resuspended in fresh medium and counted using a Cellometer auto T4 bright field cell counter (Nexcelom, Lawrence, MA) and trypan blue to distinguish viable cells. Multi-cell type tumor spheroids were grown from the mixture of three cell types: 60% malignant cells, 25% HUVEC, and 15% hMSC as described previously (42-44). Mixed cell suspensions of 42 µL were dispensed into the wells of 384-well black/clear round-bottom ULA spheroid microplates (Corning Inc., cat. 3830). Following inoculation, the microplates were transferred to an incubator (Thermo Fisher Scientific, Waltham, MA) and maintained at 37 °C and 5% CO_2_ with 95% humidity. Three days after inoculation, test agents or controls were delivered to the wells of microplates. The FDA-approved and investigational anticancer agents, prepared as 400× DMSO stock solutions, were first diluted 50-fold in media, and 6 µL were subsequently transferred to the appropriate wells of microplates containing 42 µL of cell suspension using a Tecan Freedom EVO 200 base unit (Tecan, Männedorf, Switzerland) to achieve a 1x final concentration. All anticancer agents and their combinations were tested in quadruplicate. Additionally, each microplate included a DMSO vehicle control (*n* = 16) and a cytotoxicity control (1 µM staurosporine and 3 µM gemcitabine, *n* = 20). After delivery of the test agents and controls, the microplates were returned to the incubator for 7 days. Ten days after inoculation, the assay was completed with the addition of 20 µL of CellTiter-Glo 3D (Promega, Madison, WI, cat. G9683) to each well. Next, the microplates were placed on a microplate shaker for 5 min. After 25 min of incubation at room temperature, luminescence was measured as a surrogate indicator of cell viability using a PHERAstar FSX microplate reader (BMG LABTECH, Cary, NC).

### Data Analysis

Luminescence measurements from the screen were exported as comma separated values (CSV) files and imported into custom Excel spreadsheets (Microsoft, Redmond, WA) for analysis. The raw luminescence data were evaluated for quality control, filtered for outliers, and converted to percent viability by normalizing to the DMSO (vehicle-treated) control. Concentration-response data were fit to the four-parameter logistic equation using the Solver Add-In in Excel. The Bliss independence model states that if two drugs have independent activities, then the viability for the combination is equal to the product of the viability of the two single agents (47). Synergy between two compounds is indicated by a lower observed percent viability than predicted by the Bliss independence model, whereas antagonism is indicated by a greater observed percent viability than predicted.

### Data Availability

All data are accessible via the PubChem BioAssay public database (AID 1963875; AID 1963874; AID 1963868; AID 1963873; AID 1963870; AID 1963869; AID 1963867; AID 1963866; AID 1963872; AID 1963871; AID 1963864; AID 1963865; AID 1963861; AID 1963863; AID 1963862; AID 1963860; AID 1963858; AID 1963859; AID 1963857; AID 1963856). The patient-derived cancer (PDC) cell line data used in this study can be downloaded from https://pdmr.cancer.gov/.

## RESULTS

To mimic the tumor microenvironment and facilitate cell-cell interactions, we utilized a high-throughput multi-cell type tumor spheroid model. This model is comprised of human malignant cells, HUVEC, and hMSC (42-44). Eighteen of the malignant cell lines in this study were recently derived from patients, while HOP-62 was established in the 1980s (48) (**Table S2**). Prior to investigating drug combinations (**Table S1**), the cytotoxicity of individual agents was evaluated in all the tumor spheroid models. The concentration-dependent responses from the KRAS G12C inhibitor sotorasib, the SHP2 inhibitor batoprotafib, and the SOS1 inhibitor BI-3406, across the nineteen multi-cell type tumor spheroid models are shown in **Figure 1A**. Sotorasib demonstrated relatively flat responses across the concentrations tested; however, both sotorasib and batoprotafib exhibited a degree of selectivity against tumor spheroids harboring KRAS G12C, as indicated by the normalized area under the curve heatmap (**Figure 1B**). BI-3406 showed no apparent selectivity and reached an IC_50_ in approximately half of the tumor spheroids tested. Following the initial screening of single agents, drug concentrations for combination studies were selected, based upon the clinically relevant concentration of each drug (**Table S1**). The effects of simultaneously combining agents were assessed using the Bliss independence model (47). In this model, the observed combination response is compared with a predicted combination response that is calculated from the experimentally determined response of each individual agent. Bliss independence scores nearing zero signify an additive effect, wherein the observed response closely matches the predicted outcome. A positive Bliss independence score indicates that the cell viability resulting from exposure to the combination is less than expected, demonstrating greater-than-additive cytotoxicity or synergy. A negative Bliss independence score, where the observed cell viability is greater than expected, indicates less-than-additive cytotoxicity or antagonism.

**Figure 1.**
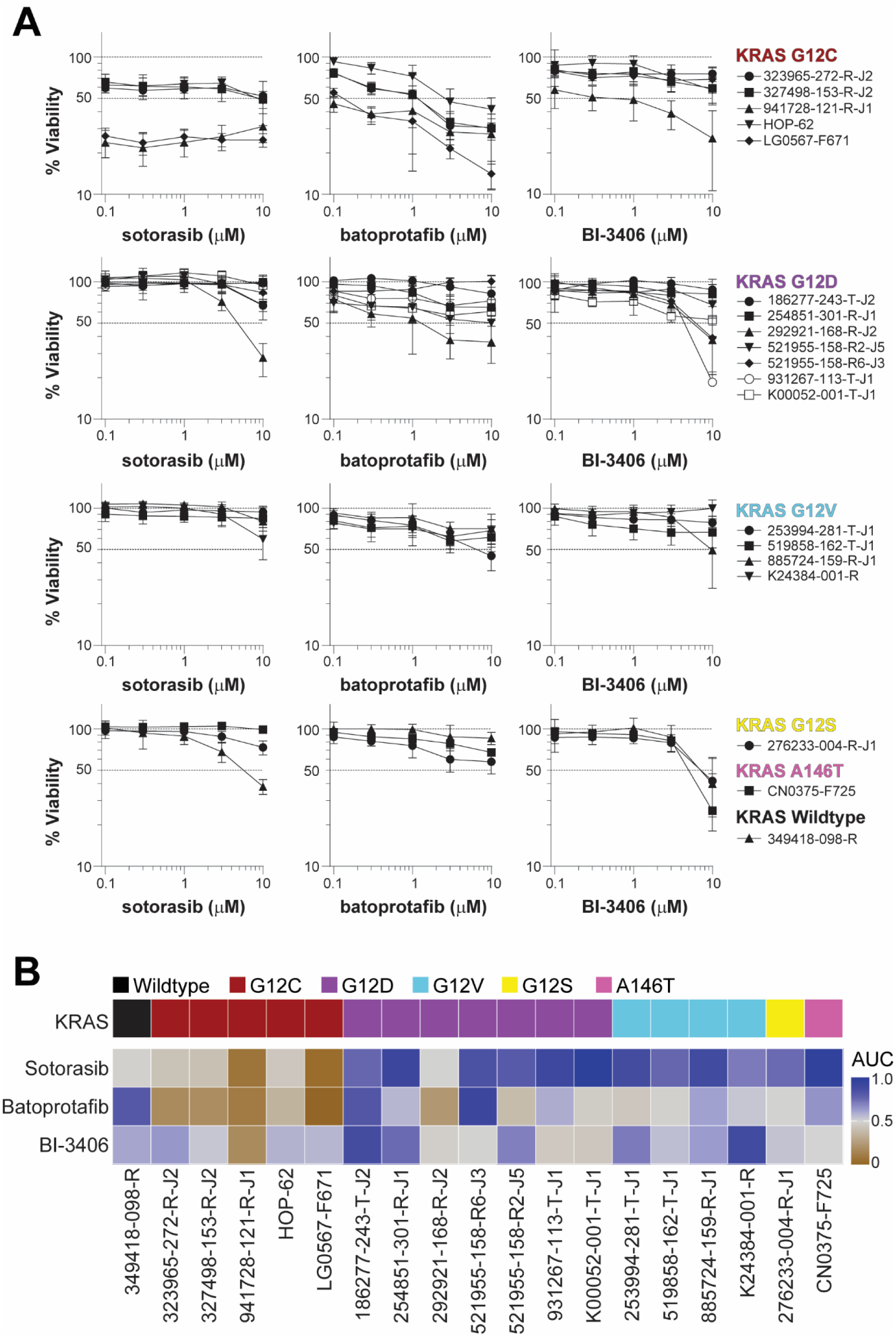
Single agent activity of sotorasib, batoprotafib, and BI-3406. (A) Concentration-response graphs from sotorasib (*left*), batoprotafib (*middle*), and BI-3406 (*right*) as single agents in all nineteen multi-cell type tumor spheroid models (mean ± SD, *n* = 4 technical replicates). (B) Heatmap of normalized area under the curve (AUC) shown in (A), with brown indicating low AUC values and blue indicating high AUC values. Annotation colors denote the KRAS genotype.

Targeting SHP2 or SOS1 along with downstream components of the RAS/MEK/ERK pathway emerged as an effective strategy. This was apparent from the combination of batoprotafib with either the MEK inhibitor trametinib or the ERK inhibitor temuterkib (LY3214996). **Figures 2A** and **3A** show concentration-response graphs and Bliss independence heatmaps from selected tumor spheroid models, with data from all models presented in **Figures S1** and **S2**. Combinations of batoprotafib with either trametinib or temuterkib showed comparable additive or synergistic effects, as indicated by the mean Bliss scores (Pearson r = 0.71, CI = 0.37 to 0.88, p = 0.0007, **Figure S3A**), across various tumor models, including a NSCLC model with wild-type KRAS and a BRAF V600E variant (349418-098-R, **Figures 2A**, **3A**, **S1**, and **S2**). Similar results were observed with the SOS1 inhibitor BI-3406 in combination with either trametinib or temuterkib. **Figures 2B** and **3B** show the concentration-response graphs and Bliss independence heatmaps from selected tumor spheroid models, with data from all models provided in **Figures S4** and **S5**. There was a weaker but statistically significant correlation between the mean Bliss scores from BI-3406 combinations with either trametinib or temuterkib (Pearson r = 0.5, CI = 0.046 to 0.78, p = 0.033, **Figure S3B**). Interestingly, statistically significant correlations were calculated between mean Bliss scores from the combinations of batoprotafib or BI-3406 with trametinib (Pearson r = 0.74, CI = 0.42 to 0.90, p = 0.0004, **Figure S3C**) or temuterkib (Pearson r = 0.85, CI = 0.63 to 0.94, p < 0.0001, **Figure S3D**), potentially attributable to the shared point at which batoprotafib and BI-3406 interfere with the RAS signaling pathway. Direct targeting of the KRAS G12C variant with sotorasib primarily resulted in additive effects when combined with trametinib or temuterkib. These additive effects were consistent across all concentrations of sotorasib, reflecting the relatively flat single agent responses over the concentrations tested. **Figures 2C** and **3C** show the concentration-response graphs and Bliss independence heatmaps for four of the five tumor spheroid models containing the KRAS G12C variant.

**Figure 2.**
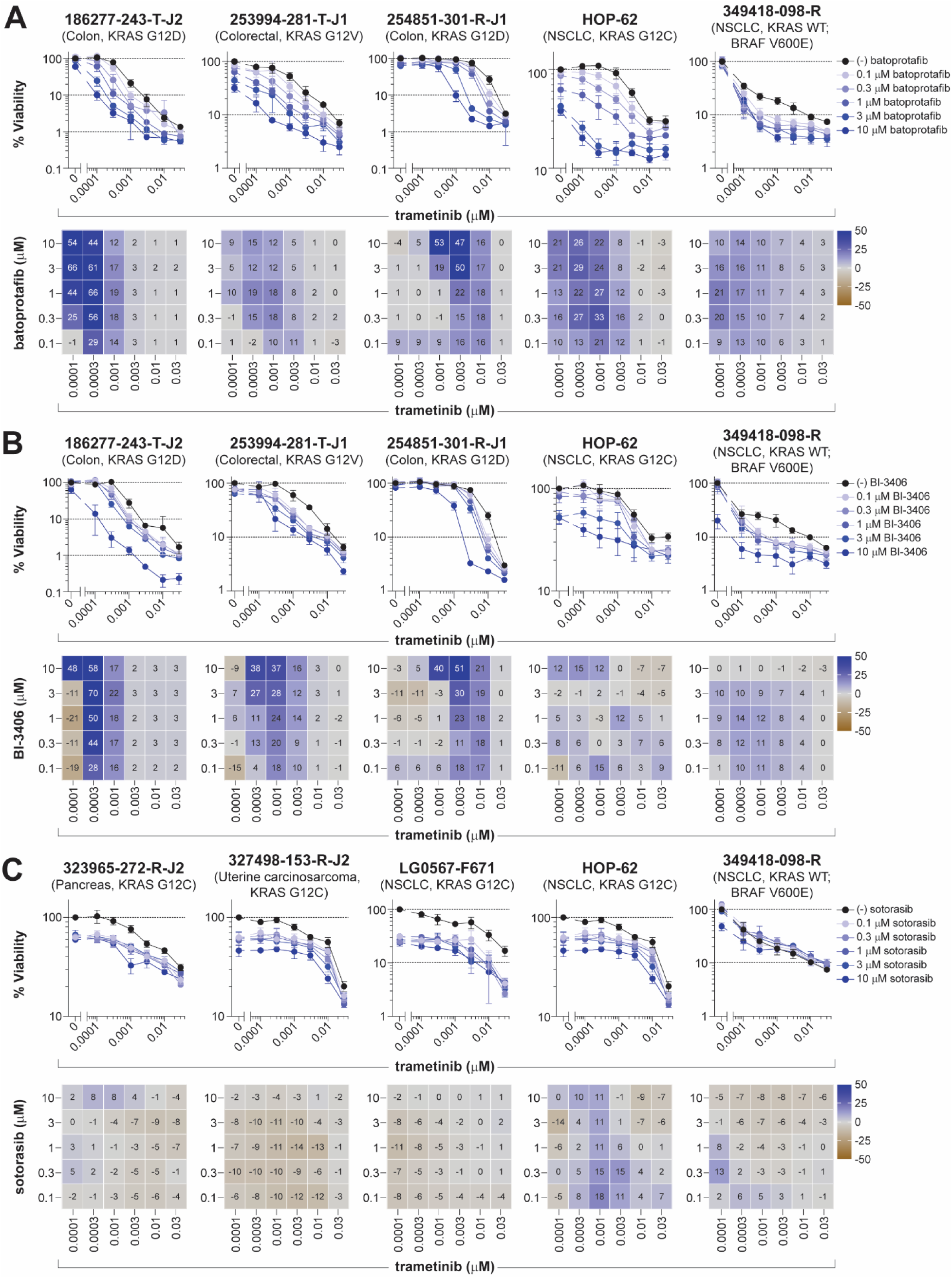
Vertical inhibition of the RAS pathway by trametinib in combination with batoprotafib, BI-3406, or sotorasib. Concentration-response graphs (*top*, mean ± SD, *n* = 4 technical replicates) from combinations of trametinib with (A) batoprotafib, (B) BI-3406, or (C) sotorasib are shown with corresponding Bliss independence scores from each combination’s concentration matrix (*bottom*, mean of *n* = 4 technical replicates) displayed numerically and as a heatmap (blue indicates synergy, gray indicates additivity, and brown indicates antagonism). The malignant cell line name, tumor type, and KRAS status are indicated above each set of graphs.

**Figure 3.**
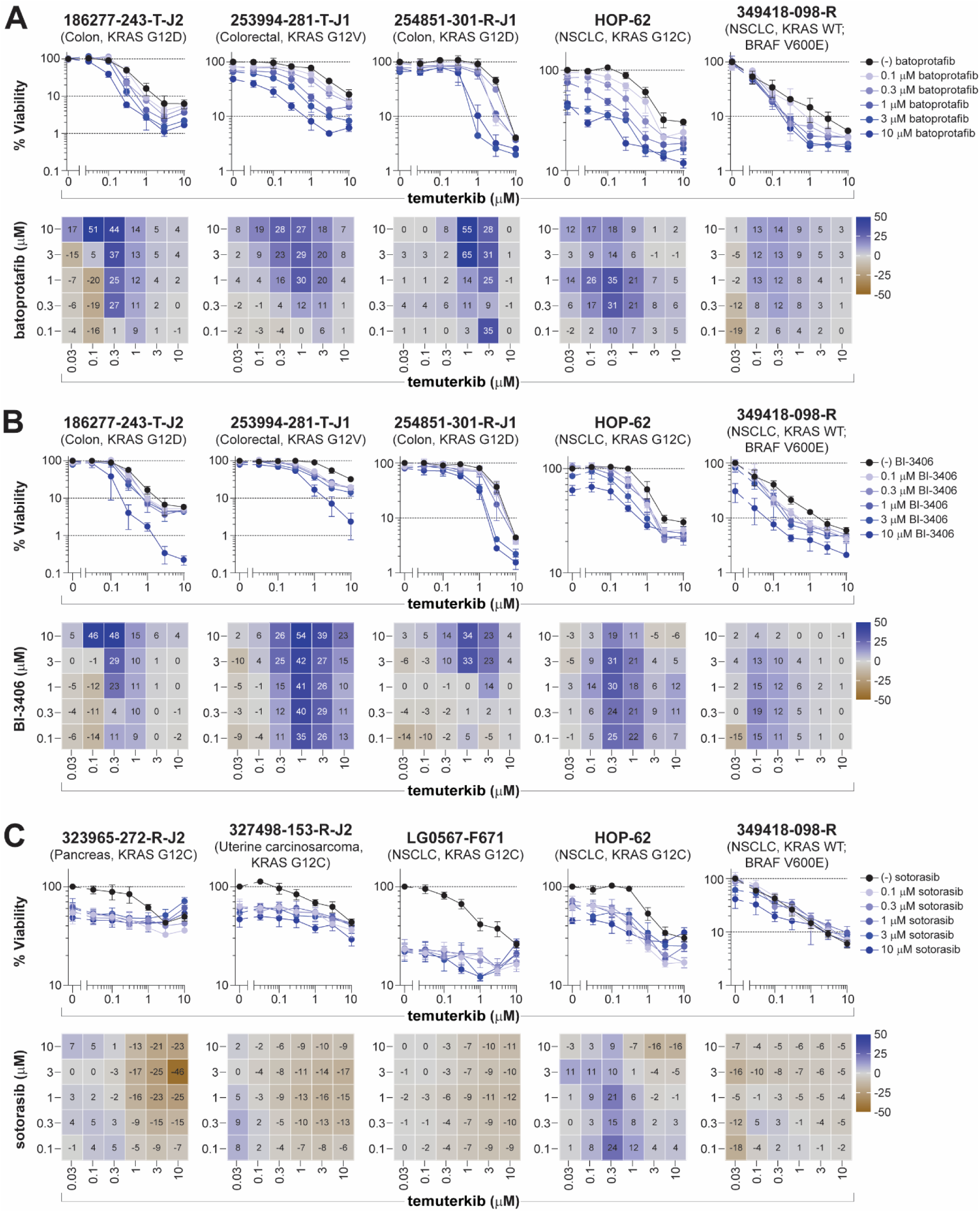
Vertical inhibition of the RAS pathway by temuterkib in combination with batoprotafib, BI-3406, or sotorasib. Concentration-response graphs (*top*, mean ± SD, *n* = 4 technical replicates) from combinations of temuterkib with (A) batoprotafib, (B) BI-3406, or (C) sotorasib are shown with corresponding Bliss independence scores from each combination’s concentration matrix (*bottom*, mean of *n* = 4 technical replicates) displayed numerically and as a heatmap (blue indicates synergy, gray indicates additivity, and brown indicates antagonism). The malignant cell line name, tumor type, and KRAS status are indicated above each set of graphs.

Inhibiting receptors upstream of the KRAS pathway with the multi-targeted receptor tyrosine kinase inhibitor nintedanib (FGFR1-3, PDGFRα/β, and VEGFR1-3) produced both additive and synergistic effects when combined with batoprotafib, BI-3406, or sotorasib. These effects were evident at the higher concentrations of each combined drug for batoprotafib and BI-3406. **Figures 4A** and **4B** show the concentration-response graphs and Bliss independence heatmaps from selected tumor spheroid models, while data from all models for the combinations of batoprotafib or BI-3406 with nintedanib are shown in **Figures S6** and **S7**, respectively. Notably, combinations of nintedanib with either batoprotafib or BI-3406 were effective against spheroids with a range of KRAS variants. Sotorasib also demonstrated both additive and synergistic effects in the tumor spheroid models harboring KRAS G12C (**Figure 4C**).

**Figure 4.**
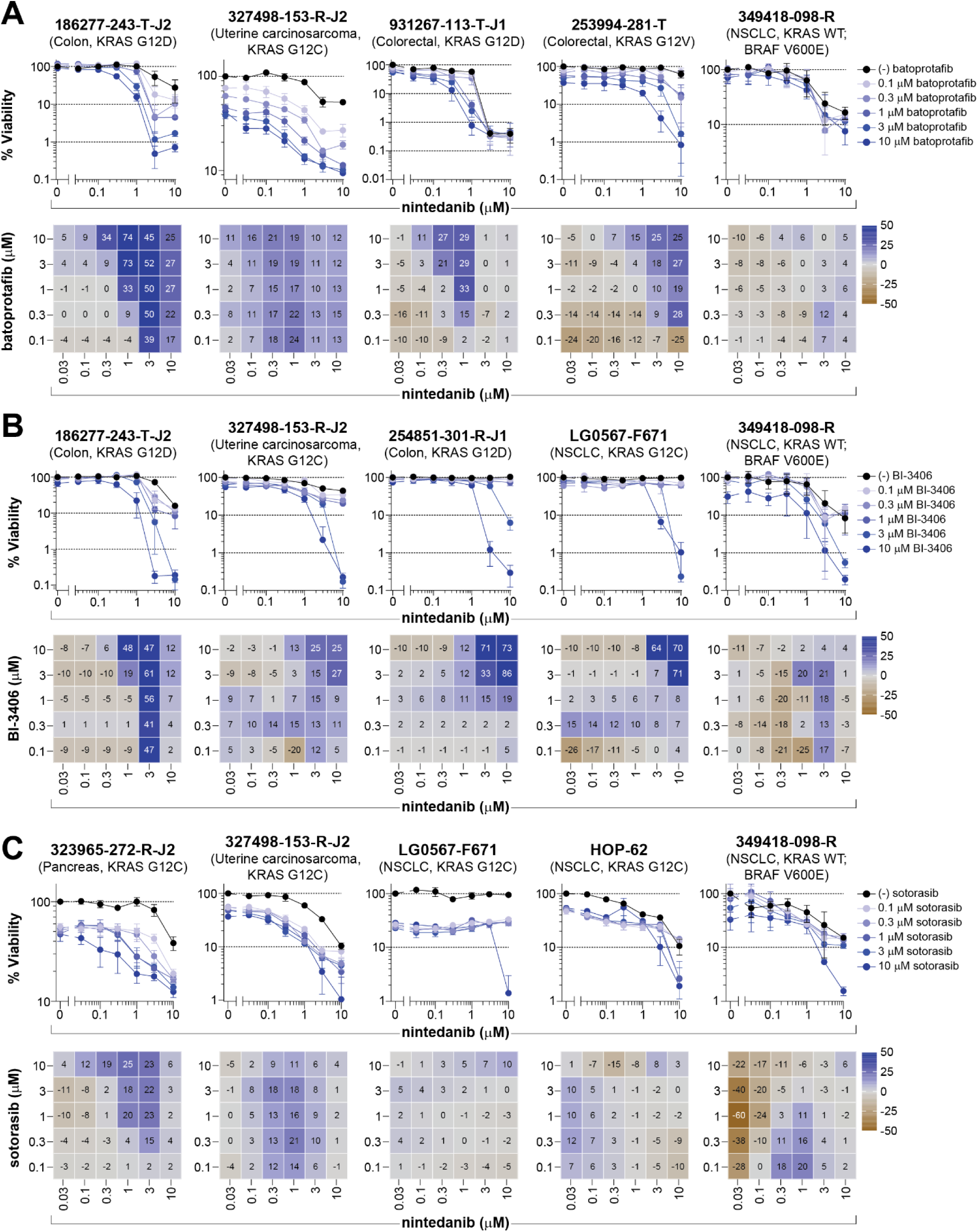
Receptor tyrosine kinase inhibition by nintenanib in combination with batoprotafib, BI-3406, or sotorasib. Concentration-response graphs (*top*, mean ± SD, *n* = 4 technical replicates) from combinations of nintedanib with (A) batoprotafib, (B) BI-3406, or (C) sotorasib are shown with corresponding Bliss independence scores from each combination’s concentration matrix (*bottom*, mean of *n* = 4 technical replicates) displayed numerically and as a heatmap (blue indicates synergy, gray indicates additivity, and brown indicates antagonism). The malignant cell line name, tumor type, and KRAS status are indicated above each set of graphs.

Targeting of both the RAS/MEK/ERK and PI3K/AKT/mTOR pathways presents another potential therapeutic strategy. Combining batoprotafib with either the mTORC1/2 inhibitor sapanisertib or the AKT inhibitor ipatasertib demonstrated both additive and synergistic activities in four models with KRAS G12C (**Figure 5A** and **6A**). No combination activity was observed in the NSCLC model with wild-type KRAS and a BRAF V600E variant (349418-098-R), and most models with variants other than G12C exhibited minimal responses to the combination of batoprotafib with sapanisertib or ipatasertib. However, synergy was observed in the patient-derived colon model 186277-243-T-J2 harboring KRAS G12D, with greater cytotoxicity observed with ipatasertib (**Figures S8** and **S9**). The combinations of BI-3406 with the PI3K/AKT/mTOR pathway inhibitors, sapanisertib or ipatasertib yielded similar results to those observed with batoprotafib, demonstrating selective activity in models with KRAS G12C and the 186277-243-T-J2 model (**Figures 5B**, **6B**, **S10** and **S11**). Sotorasib exhibited primarily additive effects in the G12C variant containing spheroids and demonstrated the greatest synergy in the NSCLC HOP-62 model (**Figures 5C** and **6C**).

**Figure 5.**
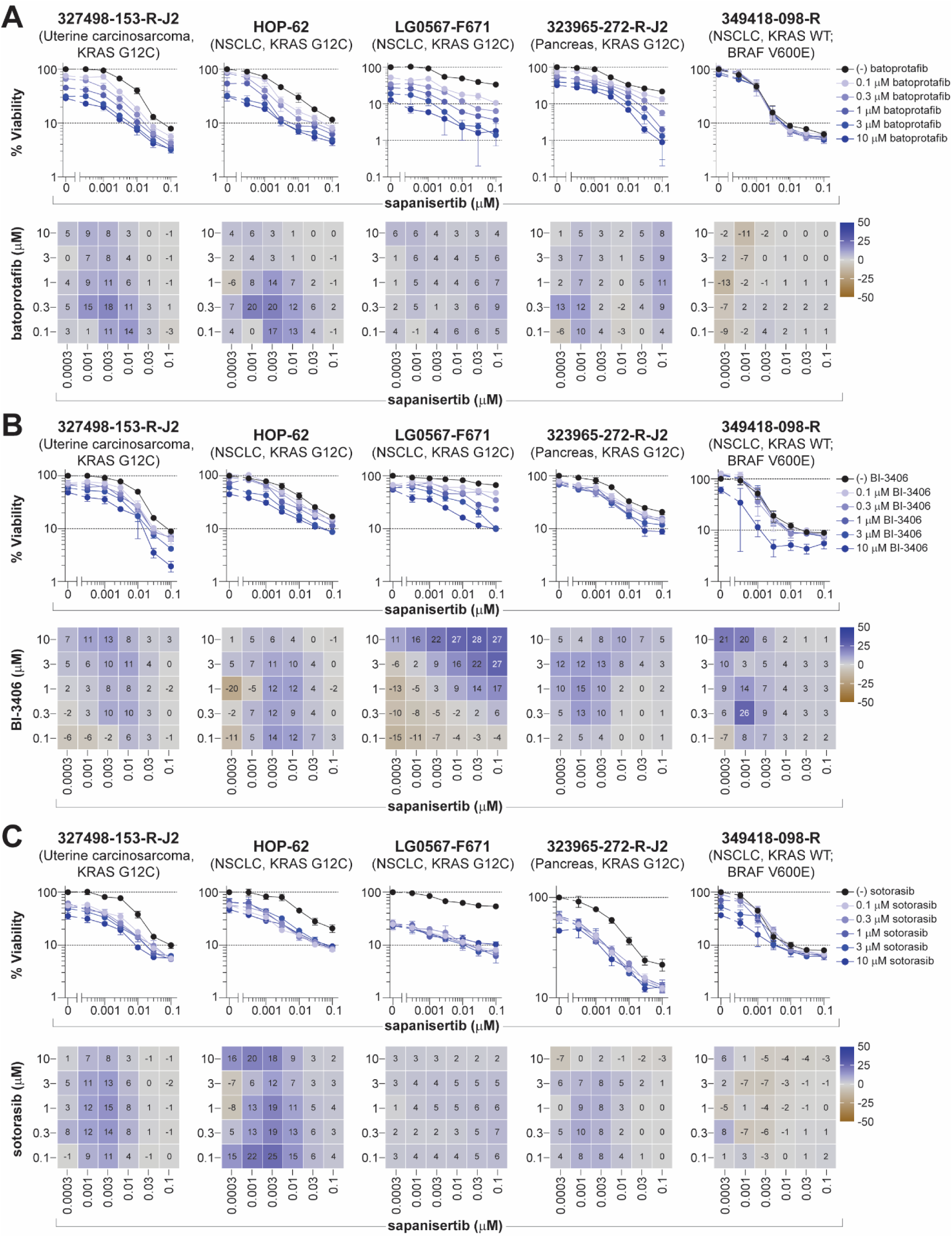
Dual pathway inhibition by sapanisertib in combination with batoprotafib, BI-3406, or sotorasib. Concentration-response graphs (*top*, mean ± SD, *n* = 4 technical replicates) from combinations of sapanisertib with (A) batoprotafib, (B) BI-3406, or (C) sotorasib are shown with corresponding Bliss independence scores from each combination’s concentration matrix (*bottom*, mean of *n* = 4 technical replicates) displayed numerically and as a heatmap (blue indicates synergy, gray indicates additivity, and brown indicates antagonism). The malignant cell line name, tumor type, and KRAS status are indicated above each set of graphs.

**Figure 6.**
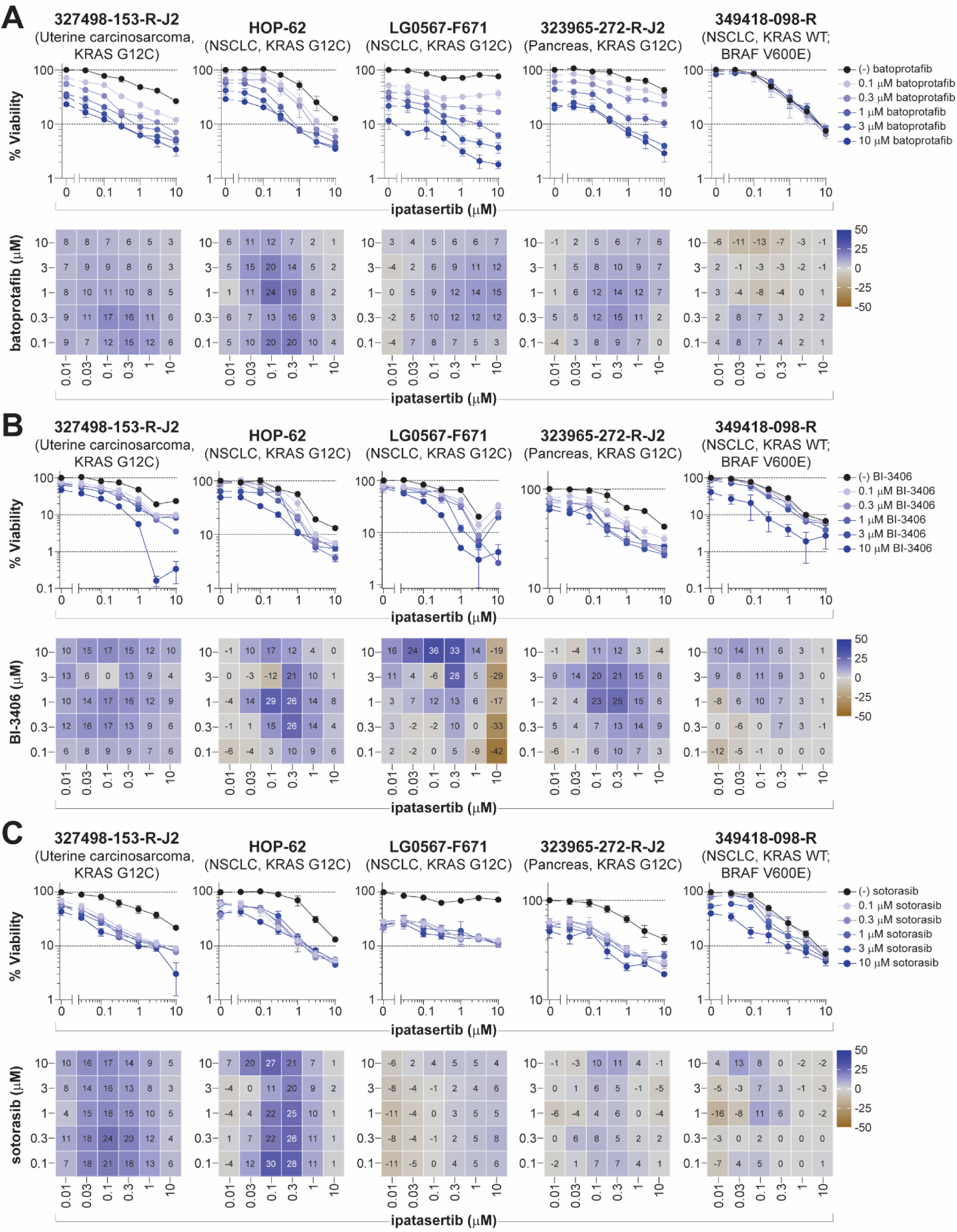
Dual pathway inhibition by ipatasertib in combination with batoprotafib, BI-3406, or sotorasib. Concentration-response graphs (*top*, mean ± SD, *n* = 4 technical replicates) from combinations of ipatasertib with (A) batoprotafib, (B) BI-3406, or (C) sotorasib are shown with corresponding Bliss independence scores from each combination’s concentration matrix (*bottom*, mean of *n* = 4 technical replicates) displayed numerically and as a heatmap (blue indicates synergy, gray indicates additivity, and brown indicates antagonism). The malignant cell line name, tumor type, and KRAS status are indicated above each set of graphs.

Combinations of batoprotafib or BI-3406 with the BCL-2 inhibitor venetoclax also demonstrated notable activity. **Figures 7A** and **7B** show concentration-response graphs and Bliss independence heatmaps from selected tumor spheroid models, while the data from all models are shown in **Figures S12** and **S13.** Greater-than-additive responses were primarily observed at higher concentrations of each agent. The combinations demonstrated additive and synergistic activity across the majority of KRAS variants but had limited impact on the 349418-098-R NSCLC model harboring wild-type KRAS and BRAF V600E. In contrast, the combination of sotorasib and venetoclax demonstrated primarily additive responses across spheroids containing the KRAS G12C variant (**Figure 7C**). Lastly, concurrent inhibition of the KRAS pathway with the combination of sotorasib and batoprotafib resulted in both additive and synergistic activity in spheroids harboring the KRAS G12C variant (**Figure 8**). Most synergy was observed in the multi-cell type tumor spheroids containing the HOP-62 NSCLC cell line, where up to one-log of cytotoxicity was achieved. Additive effects resulted in nearly 99% cytotoxicity (a two-log reduction) in the patient-derived NSCLC model LG0567-F671.

**Figure 7.**
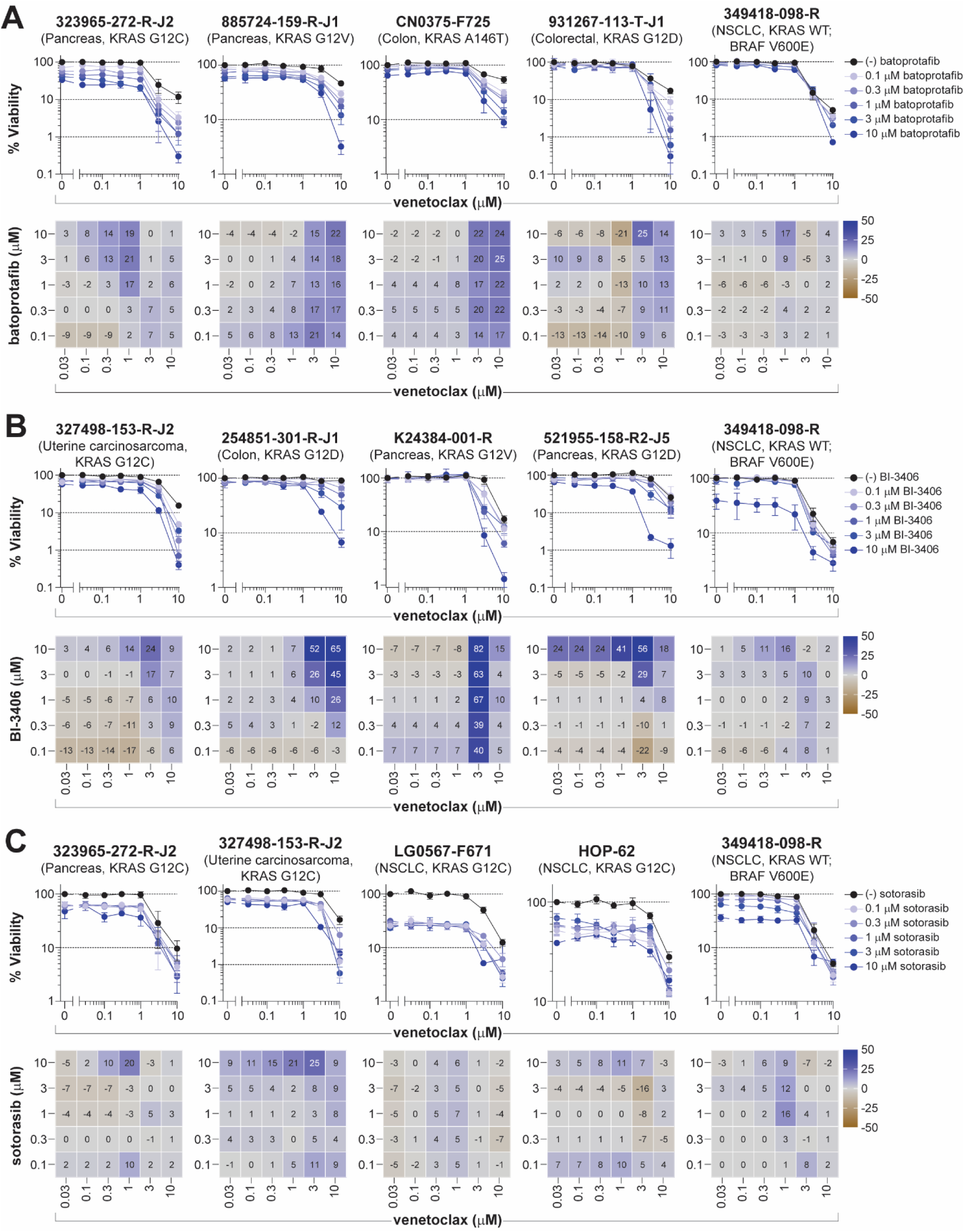
Combination of venetoclax with batoprotafib, BI-3406, or sotorasib. Concentration-response graphs (*top*, mean ± SD, *n* = 4 technical replicates) from combinations of venetoclax with (A) batoprotafib, (B) BI-3406, or (C) sotorasib are shown with corresponding Bliss independence scores from each combination’s concentration matrix (*bottom*, mean of *n* = 4 technical replicates) displayed numerically and as a heatmap (blue indicates synergy, gray indicates additivity, and brown indicates antagonism). The malignant cell line name, tumor type, and KRAS status are indicated above each set of graphs.

**Figure 8.**
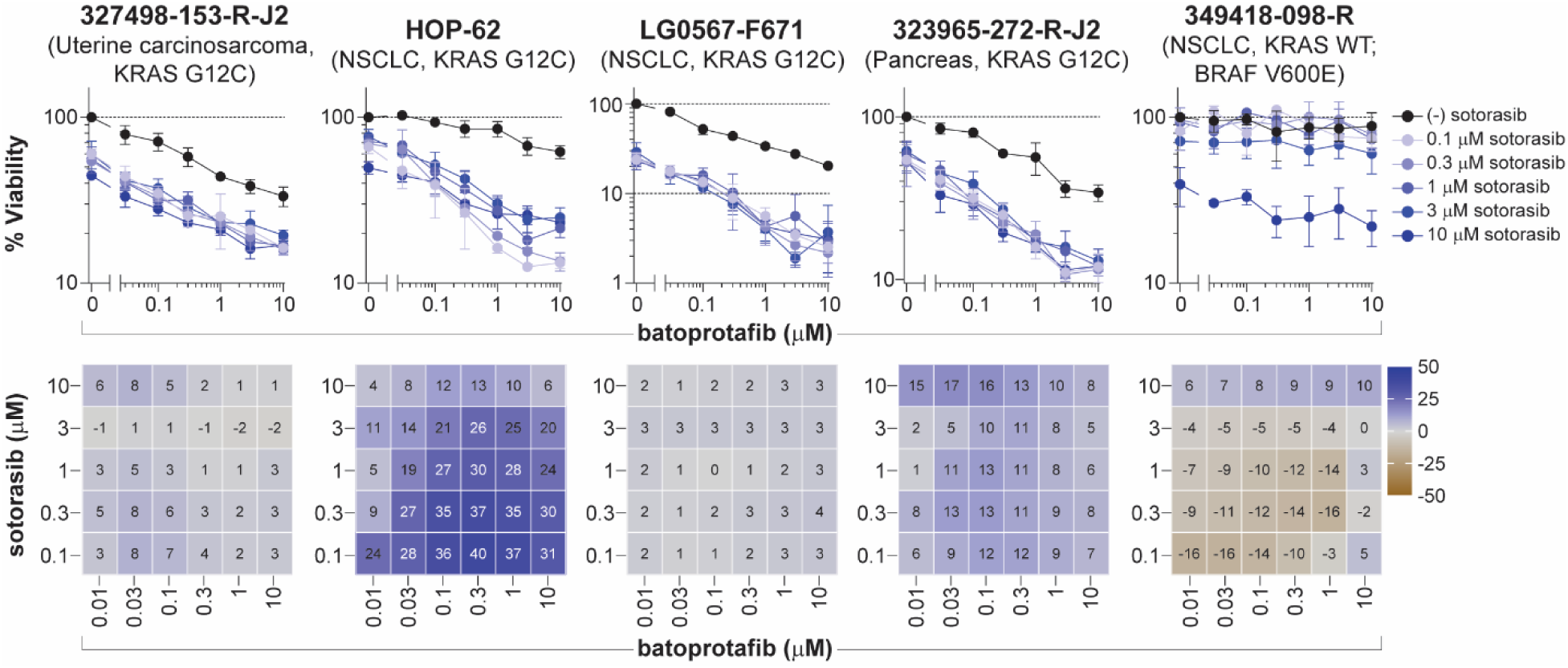
Combination of batoprotafib with sotorasib. Concentration-response graphs (*top*, mean ± SD, *n* = 4 technical replicates) from the combination of batoprotafib with sotorasib are shown with corresponding Bliss independence scores from the concentration matrix (*bottom*, mean of *n* = 4 technical replicates) displayed numerically and as a heatmap (blue indicates synergy, gray indicates additivity, and brown indicates antagonism). The malignant cell line name, tumor type, and KRAS status are indicated above each set of graphs.

## DISCUSSION

KRAS is widely recognized as one of the most frequently altered oncogenes in cancer. A recent analysis of The Cancer Genome Atlas (TCGA) database showed that KRAS variants are present in nearly 12% of all cancers, with PDAC showing the highest prevalence at over 80%, followed by CRC at nearly 40%, and approximately 20% in patients with NSCLC (49). The landmark FDA approval of sotorasib for treatment of NSCLC bearing the KRAS G12C variant proved that exquisite targeting of malignant cells can have a therapeutic benefit (27). Approximately one year later, the KRAS G12C inhibitor adagrasib received accelerated approval by the FDA for the same indication (50). A range of additional KRAS G12C inhibitors are currently under development and include: olomorasib (LY3537982), divarasib (GDC-6036), garsorasib (D-1553), HBI-2438, opnurasib (JDQ443), glecirasib (JAB-21822), HS-10370, IBI351 (GFH925), BI 1823911, JNJ-74699157, ARS-853, ARS-1620, ASP2453, and ERAS-3490 (6). Inhibitors targeting other KRAS variants, including G12D, G12S, and G12R are also under development (51-53), with several G12D-selective inhibitors now progressing through clinical trials (54). Beyond directly targeting specific KRAS variants, the development of pan-KRAS(ON) inhibitors that target the full spectrum of KRAS variants shows promise for effectively treating a range of cancers (55-57). It has recently been noted that sotorasib is an effective inhibitor not only of KRAS G12C but also HRAS G12C and NRAS G12C, expanding the range and number of tumors where sotorasib might be therapeutically important (58). Although sotorasib treatment resulted in significant responses in the targeted patient population, durable complete responses were not achieved. Improved treatment for patients with KRAS G12C bearing NSCLC will likely come from combination therapeutic regimens including immunotherapy, radiation therapy, and chemotherapy.

There have also been efforts to suppress RAS pathway activity by targeting signaling upstream of RAS. Two such efforts include targeting SHP2 and SOS1, both of which disrupt the RAS nucleotide exchange process (59). Clinical trials of SHP2 and SOS1 inhibitors as a monotherapy have demonstrated limited efficacy, so drug combinations are under clinical evaluation (60-62). SHP2 has been shown to regulate immunological checkpoints and inhibit antitumor immune responses (63,64). Consequently, several clinical trials are evaluating the combination of SHP2 inhibitors with immunotherapy (clinicaltrials.gov; NCT05375084, NCT05505877, NCT04000529).

In this study, the allosteric SHP2 inhibitor batoprotafib, the SOS1 inhibitor BI-3406, and the KRAS G12C allele-specific inhibitor sotorasib were examined in combination with a range of targeted small molecule agents (**Table S1**). Potential approaches explored for combination chemotherapy regimens included targeting RTKs (erdafitinib, nintedanib, cabozantinib), mitogen-activated protein kinases (trametinib, temuterkib), PI3K/AKT/mTOR signaling pathway intermediates (ipatasertib, sapanisertib) or complimentary targets such as BCL-2 (venetoclax) or PARP (olaparib, talazoparib). Single agents and combinations were evaluated in multi-cell type tumor spheroid models grown from nineteen malignant cell lines derived from cancers with a range of oncogenic KRAS variants (**Table S2**).

As single agents, sotorasib and batoprotafib demonstrated selectivity towards tumor spheroids harboring KRAS G12C, whereas BI-3406 elicited responses in a broader range of models (**Figure 1A** and **1B**). However, previous studies have shown that cell lines expressing KRAS G12C or G12A demonstrated selective sensitivity to the allosteric SHP2 inhibitor RMC-4550 (65). The activity of SHP2 inhibitors correlates with KRAS GTPase activity (66,67). This can partly be attributed to the differential impact of KRAS variants on GTPase activity or GAP-mediated hydrolysis, wherein G12C compared to other variants, demonstrated an intrinsic GTP hydrolysis rate nearly equivalent to that of wild-type KRAS (68). Conversely, SOS1 inhibitors disrupt the protein-protein interaction between SOS1 and KRAS by binding the SOS1 catalytic site, preventing nucleotide exchange and KRAS GTP loading. This mechanism suggests their potential efficacy against a broad range of KRAS variants (34,69).

Efforts to target downstream effectors of RAS have involved inhibitors of BRAF, MEK, and ERK (70-72). However, those strategies have faced challenges from compensatory feedback mechanisms, which can lead to reactivation of multiple RTKs and the emergence of therapeutic resistance (73). Both SOS1 and SHP2 function as proximal signaling intermediates for RTKs upstream of RAS and are essential for RTK-dependent RAS activation and subsequent pathway feedback mechanisms. Among the drug combinations evaluated in this study, combination effects were frequently observed from concurrently blocking a downstream effector and an upstream activator of KRAS, irrespective of the KRAS status. For instance, vertical pathway inhibition with the SHP2 inhibitor batoprotafib in combination with either the MEK inhibitor trametinib or the ERK inhibitor temuterkib led to additive and/or synergistic responses (**Figure 2A** and **3A**). Similar results were observed with the SOS1 inhibitor BI-3406 (**Figure 2B** and **3B**). The consistency of these effects across the tumor spheroid models is indicated by correlations in mean Bliss scores (**Figure S3**) and reflects the similar point at which SHP2 and SOS1 inhibitors disrupt the RAS signaling pathway. Several clinical trials are underway to evaluate combinations of a MEK inhibitor with the SOS1 inhibitor BI 1701963 (closely related to BI-3406) or a SHP2 inhibitor (clinicaltrials.gov; NCT03989115, NCT04292119, NCT04800822 (74), and NCT04111458 (75)). Combinations of a SHP2 inhibitor with an ERK inhibitor are also undergoing evaluation in the clinic (clinicaltrials.gov; NCT04866134 and NCT04916236 (76)). Although the combination activity of sotorasib with either trametinib or temuterkib was modest compared to the more pronounced activity observed with batoprotafib or BI-3406 (**Figure 2C** and **3C**), antitumor activity was observed for the combination of sotorasib and trametinib, including responses in patients previously treated with KRAS G12C inhibitors (77). Ongoing clinical trials continue to explore downstream targets of RAS in combination with KRAS G12C inhibitors.

Both SHP2 and SOS1 inhibitors exhibited cytotoxicity in RTK-driven human cancer cells, suggesting that inhibitors targeting oncogenic RTKs upstream of RAS might sensitize cells to SHP2 or SOS1 inhibition (38,78). The combined pharmacological inhibition of SHP2 or SOS1 with EGFR-variant specific inhibitors, such as nazartinib and osimertinib, produced synergistic effects in both NSCLC cell lines and patient-derived xenografts with an EGFR-variant (78-80). In addition, blocking SHP2 in conjunction with an FGFR-targeted kinase inhibitor synergistically inhibited the growth of metastatic breast cancer cells and suppressed metastatic progression *in vivo* (81). In the multi-cell type tumor spheroids, combinations of batoprotafib or BI-3406 with nintedanib, a multi-targeted receptor tyrosine kinase inhibitor, demonstrated additive and synergistic responses at the higher concentrations of each agent and across KRAS variants (**Figure 4A** and **4B**). Several clinical trials are currently assessing the combination of SHP2 inhibitors with RTK inhibitors, including anlotinib, a multi-targeted receptor tyrosine kinase inhibitor (clinicaltrials.gov; NCT05715398), as well as two EGFR inhibitors, nazertinib (clinicaltrials.gov; NCT03114319) and osimertinib (clinicaltrials.gov; NCT03989115). The most substantial combination effects for sotorasib with nintedanib were observed in the KRAS G12C containing pancreatic 323965-272-R-J2 and uterine carcinosarcoma 327498-153-R-J2 tumor spheroid models (**Figure 4C**). Clinical results from combining sotorasib with the EGFR inhibitor afatinib in NSCLC patients demonstrated overall response rates (ORR) of 20% and 34.8%, with disease control rates (DCR) of 70% and 73.9% across two dose cohorts (82). Additionally, combining KRAS G12C inhibitors with anti-EGFR monoclonal antibodies has shown encouraging antitumor activity compared to the standard of care in colorectal cancer (83-86).

The RAS/MEK/ERK and PI3K/AKT/mTOR pathways continue to be pivotal targets for anticancer therapies (87,88). In preclinical studies, simultaneous inhibition of both pathways demonstrated greater efficacy than targeting a single pathway. However, translating this approach to human patients has been challenging due to dose-limiting toxicities and the difficulty in establishing a therapeutic window (89-91). Combination effects were apparent in the multi-cell type tumor spheroids when pairing batoprotafib with the PI3K/AKT/mTOR pathway inhibitors sapanisertib or ipatasertib (**Figure 5A** and **6A**). Notably, these combinations displayed relatively selective activity towards tumor spheroids harboring KRAS G12C. The combination of sapanisertib or ipatasertib with BI-3406 showed similar activity and selectivity (**Figure 5B** and **6B**). Similar outcomes were observed with the concurrent inhibition of SHP2 and the PI3K/AKT/mTOR pathway in *in vitro* and *in vivo* models of liver, ovarian, and metastatic breast cancer (92-94). Dual pathway inhibition with sotorasib combined with either sapanisertib or ipatasertib also exhibited combination activity (**Figure 5C** and **6C**). Targeting the PI3K/AKT/mTOR pathway has been shown to be effective when combined with KRAS G12C inhibitors (22,41,95). For example, sotorasib in combination with an mTORC1/2 inhibitor led to sustained inhibition of tumor growth and metastasis in a mouse model of pancreatic ductal adenocarcinoma (96). There are two clinical trials investigating the combination of KRAS G12C inhibitors with mTOR inhibitors in solid tumors harboring KRAS G12C (clinicaltrials.gov; NCT05840510 and NCT04185883).

Combination effects were also observed in various spheroid models when batoprotafib, BI-3406 or sotorasib was combined with the BCL-2 inhibitor venetoclax (**Figure 7**). While responses from the combination of sotorasib with venetoclax were primarily additive in models harboring KRAS G12C, greater-than-additive responses were observed from combinations of venetoclax with batoprotafib and BI-3406. Recently, synergy between the allosteric SHP2 inhibitor RMC-4550 and venetoclax was observed in acute myeloid leukemia (AML) models with alterations in *FLT3* and *KIT*, indicating that SHP2 inhibition enhances cellular dependence on BCL-2 (97). The combination of a SOS1 inhibitor with venetoclax improved efficacy in an AML cell line and patient-derived xenograft models compared with either agent alone (98). In clinical settings, BCL-2 inhibitors seem to have less efficacy against solid tumors compared to hematological cancers when used as single-agent therapy (99). Results from the multi-cell type tumor spheroid models suggest that KRAS, SHP2 or SOS1 inhibitors could enhance the effectiveness of BCL-2 inhibitors against solid tumors; however, hematologic toxicity might be a limitation.

The combination of sotorasib with batoprotafib exhibited both additive and synergistic responses in KRAS G12C tumor spheroids (**Figure 8**). Mechanistically, inhibiting SHP2 or SOS1 reduces the GDP–GTP exchange rate and increases the occupancy of the KRAS G12C-GDP state, which is targeted by some KRAS G12C inhibitors (12). Multiple clinical trials are exploring combinations of KRAS G12C inhibitors with SHP2 or SOS1 inhibitors (100-103). Recent clinical results suggest promising outcomes for this combination strategy. For instance, combination of the KRAS G12C inhibitor glecirasib with the SHP2 inhibitor JAB-3312 demonstrated an ORR of 50% (14/28) and a DCR of 100% in patients with KRAS G12C inhibitor-naive NSCLC (104).

Altogether, the findings in these patient-derived tumor spheroid models confirm and broaden the understanding of drug combinations with inhibitors of KRAS G12C, SHP2 and SOS1. Many of the combination strategies evaluated in this study are undergoing investigation in clinical trials. Continued research will be essential to validate the efficacy of these combination strategies and their potential for cancer therapy.

