## Supplemental Material for "Combinations of RAS pathway inhibitors with targeted agents are active in spheroids of patient-derived cells with oncogenic KRAS variants from multiple cancer types"

**Table S1**. Drugs and investigational agents used in this study. The agents included RAS pathway inhibitors (*top*) and molecular targeted combination drugs (*bottom*). If available at the time of this study, the clinical $C_{max}$ is listed.

| **RAS Pathway Inhibitors** | **Clinical $C_{max}$** | **Molecular Target** |
|---|---|---|
| sotorasib (AMG 510) | 9.61 µM | KRAS G12C |
| MRTX1257 | *NA* | KRAS G12C |
| batoprotafib (TNO155) | *NA* | SHP2 |
| BI-3406 | *NA* | SOS1 |
| BAY-293 | *NA* | SOS1 |
| **Combination Drugs** | **Clinical $C_{max}$** | **Molecular Target** |
| elimusertib (BAY 1895344) | *NA* | ATR |
| molibresib (GSK525762) | *NA* | BET bromodomain |
| temuterkib (LY3214996) | *NA* | ERK |
| venetoclax | 4.48 µM | BCL-2 |
| alisertib | *NA* | Aurora A kinase |
| olaparib | 13.1 µM | PARP |
| talazoparib | 0.043 µM | PARP |
| erdafitinib | 3.1 µM | FGFR |
| ipatasertib | *NA* | AKT1/2/3 |
| cabozantinib | 4.61 µM | cMET, VEGFR, cKIT |
| nintedanib | 4 µM | PDGFR, FGFR, VEGFR |
| abemaciclib | 0.588 µM | CDK 4/6 |
| docetaxel | 5.47 µM | Tubulin stabilizer |
| trametinib | 0.021 µM | MEK 1/2 |
| sapanisertib | 0.96 µM | mTORC1/2 |
| batoprotafib (TNO155) | *NA* | SHP2 |

*NA*, clinical $C_{max}$ unknown, highest concentration tested was 10 µM.

**Table S2**. The malignant cell lines grown as multi-cell type tumor spheroids for this study. The names of both patient-derived and established cell lines are listed along with the tumor type they were derived from and the KRAS status.

| Cell Line | Tumor Type | KRAS Status |
|---|---|---|
| 186277-243-T-J2 | Colon cancer | KRAS G12D |
| 254851-301-R-J1 | Colon cancer | KRAS G12D* |
| 276233-004-R-J1 | Colon cancer | KRAS G12S |
| 519858-162-T-J1 | Colon cancer | KRAS G12V |
| CN0375-F725 | Colon cancer | KRAS A146T |
| 931267-113-T-J1 | Colorectal cancer | KRAS G12D |
| 253994-281-T-J1 | Colorectal cancer | KRAS G12V |
| LG0567-F671 | Non-small cell lung cancer | KRAS G12C |
| 941728-121-R-J1 | Non-small cell lung cancer | KRAS G12C |
| K00052-001-T-J1 | Non-small cell lung cancer | KRAS G12D |
| 349418-098-R | Non-small cell lung cancer | KRAS WT (BRAF V600E) |
| HOP-62 | Non-small cell lung cancer | KRAS G12C |
| K24384-001-R | Pancreatic cancer | KRAS G12V |
| 292921-168-R-J2 | Pancreatic cancer | KRAS G12D* |
| 323965-272-R-J2 | Pancreatic cancer | KRAS G12C |
| 885724-159-R-J1 | Pancreatic cancer | KRAS G12V |
| 521955-158-R2-J5 | Pancreatic cancer | KRAS G12D |
| 521955-158-R6-J3 | Pancreatic cancer | KRAS G12D |
| 327498-153-R-J2 | Uterine carcinosarcoma | KRAS G12C |

All data were obtained from the NCI Patient-Derived Models Repository (https://pdmr.cancer.gov) or cBioPortal (https://www.cbioportal.org/).

*From the OncoKB Gene Panel of the autologous organoid model (PDC data were not available).

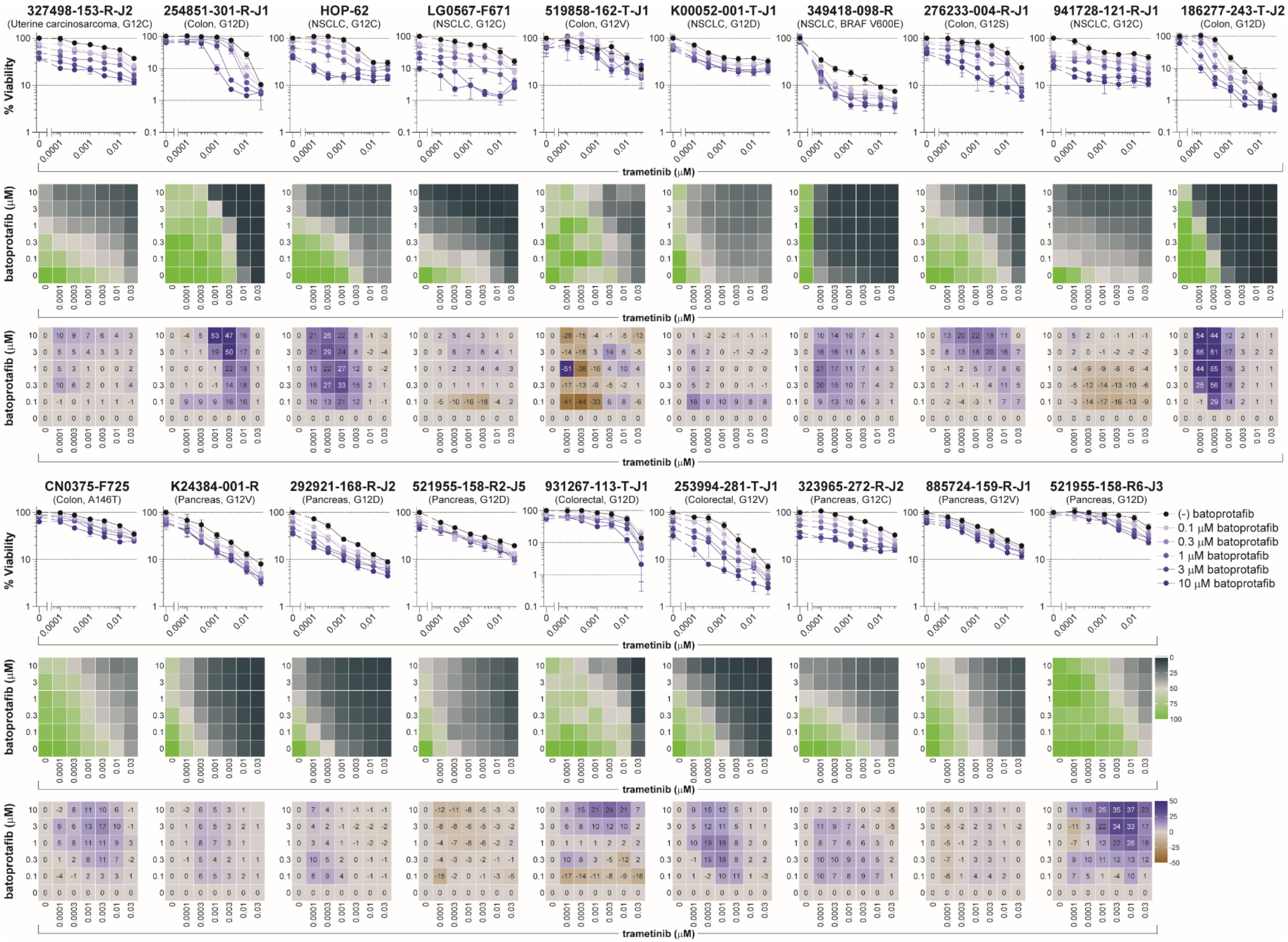

**Figure S1**. **Combination activity for batoprotafib with trametinib in multi-cell type tumor spheroids**. Concentration-response graphs (*top*, mean ± SD, $n$ = 4 technical replicates), % viability across the combination's concentration matrix (*middle*, mean of $n$ = 4 technical replicates) displayed as a heatmap (green indicates high cell viability and black indicates low cell viability), and Bliss independence scores across the combination's concentration matrix (*bottom*, mean of $n$ = 4 technical replicates) displayed numerically and as a heatmap (blue indicates synergy, gray indicates additivity, and brown indicates antagonism) are shown for nineteen malignant cell lines grown as multi-cell type tumor spheroids and exposed to batoprotafib in combination with trametinib. The malignant cell line name, tumor type, and KRAS status are indicated above each set of graphs.

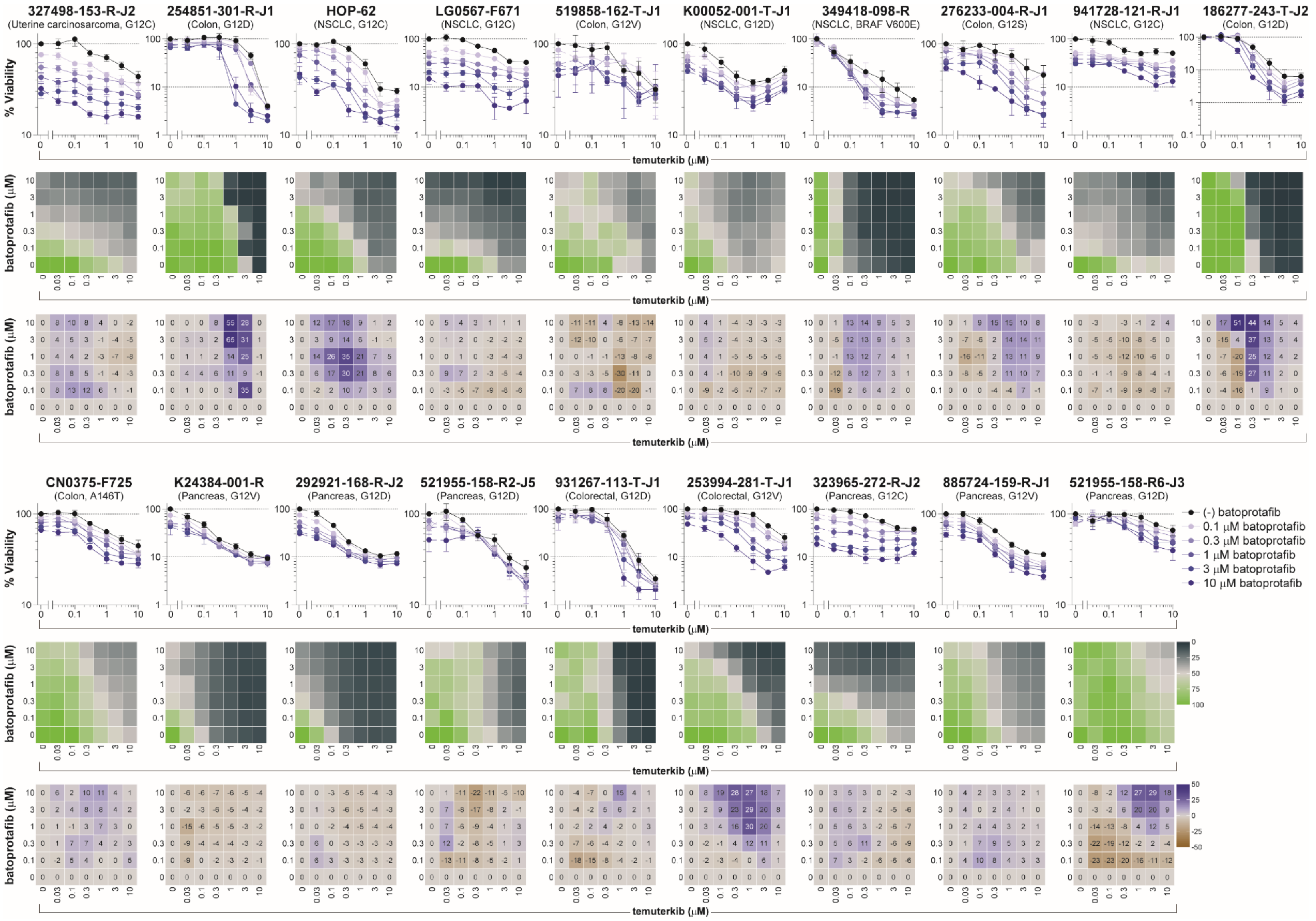


**Figure S2**. **Combination activity for batoprotafib with temuterkib in multi-cell type tumor spheroids**. Concentration-response graphs (*top*, mean ± SD, *n* = 4 technical replicates), % viability across the combination's concentration matrix (*middle*, mean of *n* = 4 technical replicates) displayed as a heatmap (green indicates high cell viability and black indicates low cell viability), and Bliss independence scores across the combination's concentration matrix (*bottom*, mean of *n* = 4 technical replicates) displayed numerically and as a heatmap (blue indicates synergy, gray indicates additivity, and brown indicates antagonism) are shown for nineteen malignant cell lines grown as multi-cell type tumor spheroids and exposed to batoprotafib in combination with temuterkib. The malignant cell line name, tumor type, and KRAS status are indicated above each set of graphs.

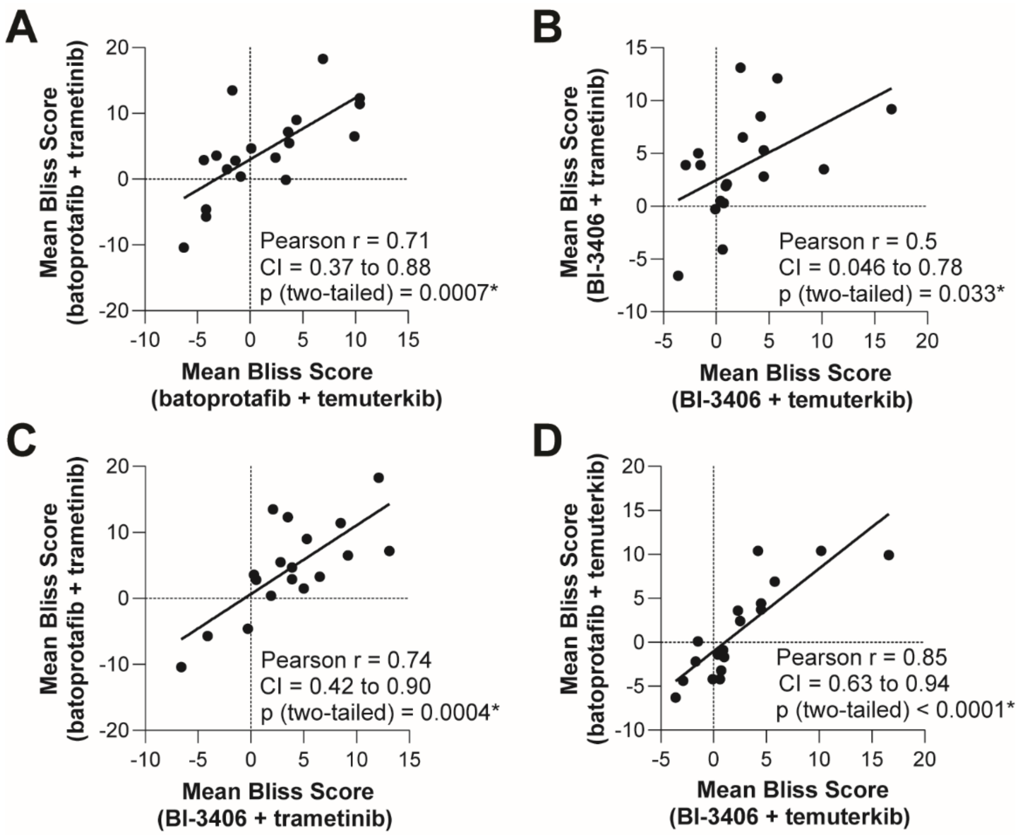


**Figure S3**. **Mean Bliss score correlations for vertical inhibition of the KRAS pathway by batoprotafib or BI-3406 in combination with trametinib or temuterkib**. Scatter plots depict significant correlations between the mean Bliss scores from combinations of (A) batoprotafib with trametinib or temuterkib, (B) BI-3406 with trametinib or temuterkib, (C) trametinib with batoprotafib or BI-3406, and (D) temuterkib with batoprotafib or BI3406. Pearson correlation coefficients (r), confidence intervals (CI), and p-values are shown, with statistical significance indicated by an asterisk (*).

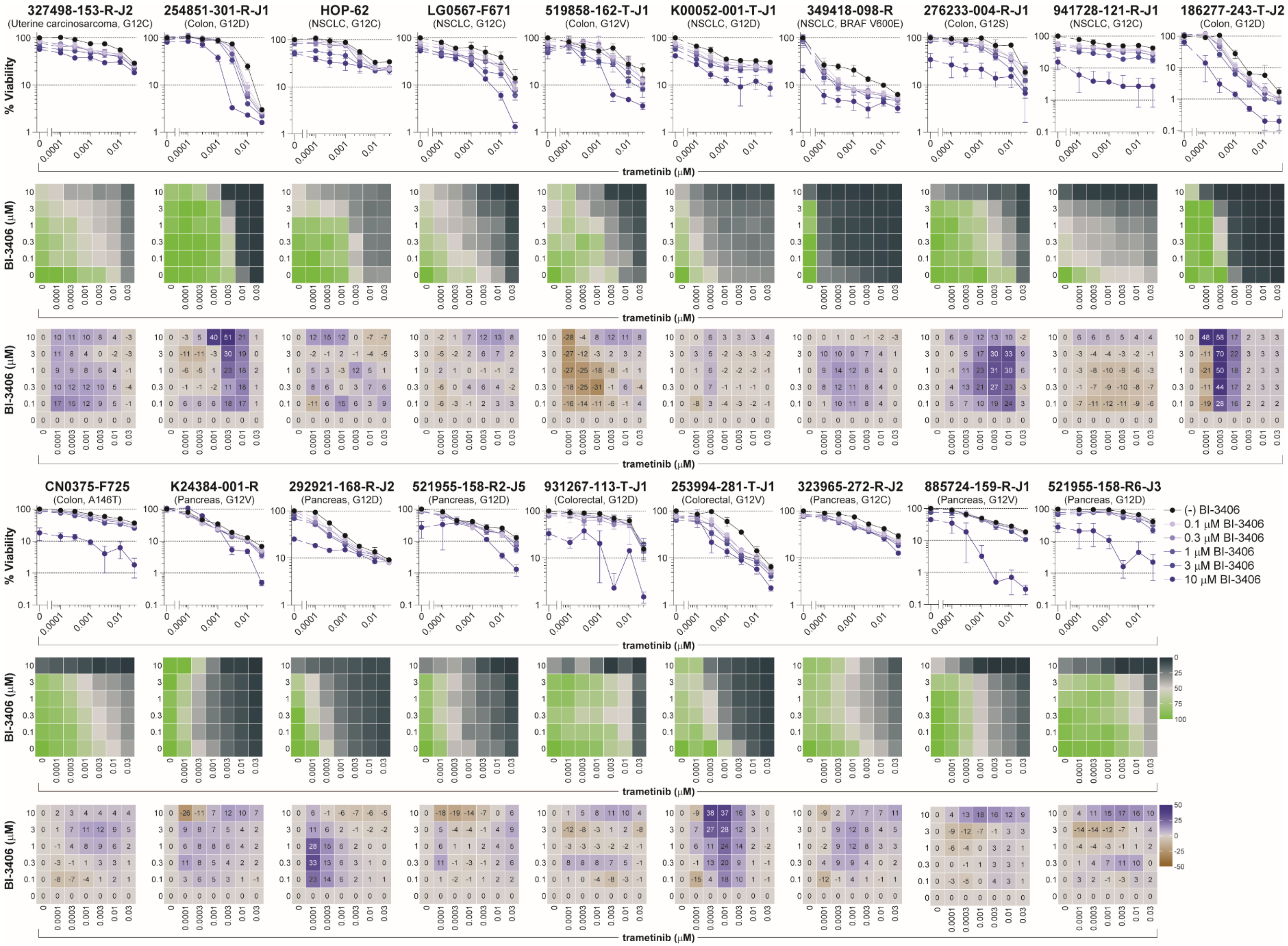


**Figure S4**. **Combination activity for BI-3406 with trametinib in multi-cell type tumor spheroids**. Concentration-response graphs (*top*, mean ± SD, *n* = 4 technical replicates), % viability across the combination's concentration matrix (*middle*, mean of *n* = 4 technical replicates) displayed as a heatmap (green indicates high cell viability and black indicates low cell viability), and Bliss independence scores across the combination's concentration matrix (*bottom*, mean of *n* = 4 technical replicates) displayed numerically and as a heatmap (blue indicates synergy, gray indicates additivity, and brown indicates antagonism) are shown for nineteen malignant cell lines grown as multi-cell type tumor spheroids and exposed to BI-3406 in combination with trametinib. The malignant cell line name, tumor type, and KRAS status are indicated above each set of graphs.

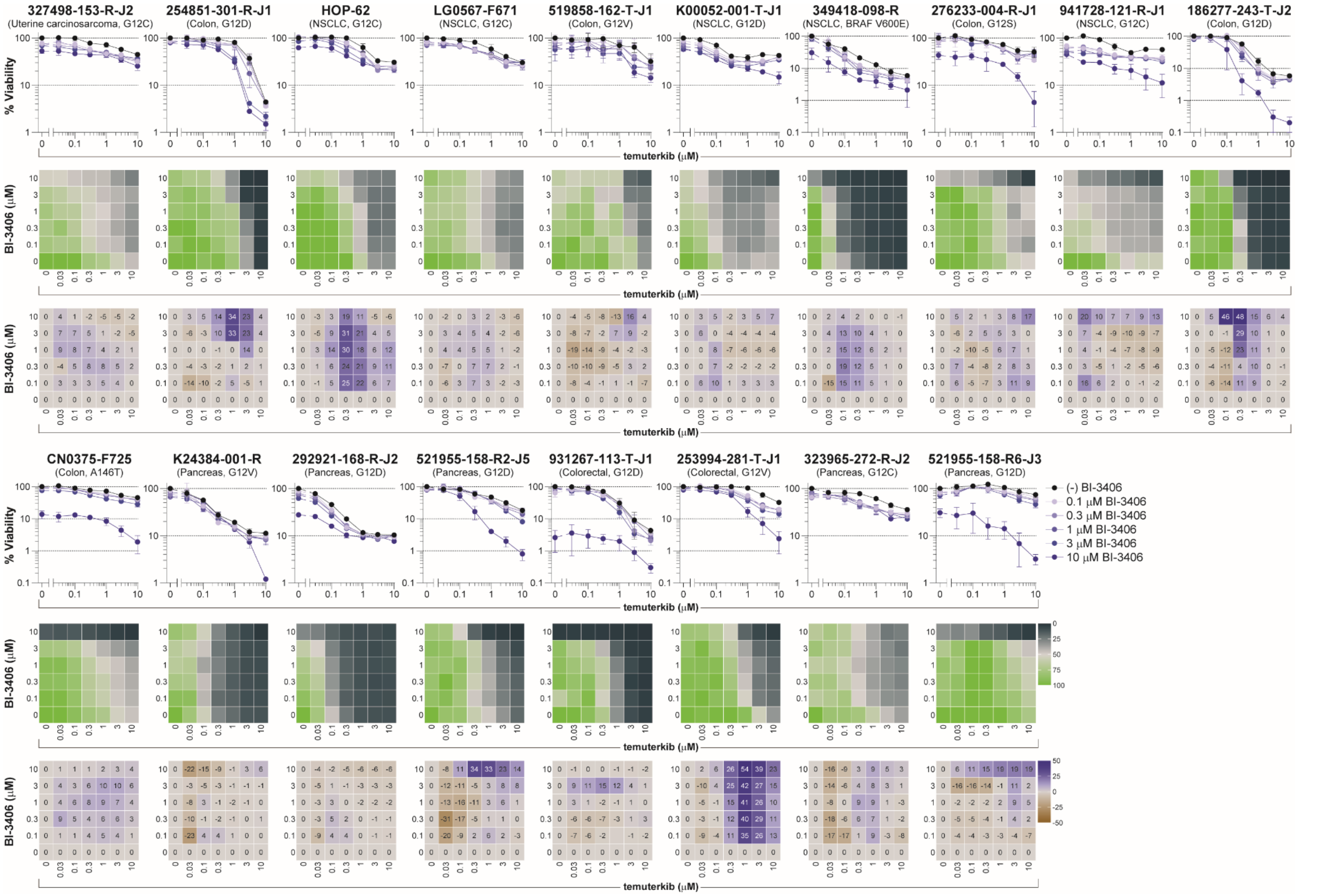


**Figure S5**. **Combination activity for BI-3406 with temuterkib in multi-cell type tumor spheroids**. Concentration-response graphs (*top*, mean ± SD, *n* = 4 technical replicates), % viability across the combination's concentration matrix (*middle*, mean of *n* = 4 technical replicates) displayed as a heatmap (green indicates high cell viability and black indicates low cell viability), and Bliss independence scores across the combination's concentration matrix (*bottom*, mean of *n* = 4 technical replicates) displayed numerically and as a heatmap (blue indicates synergy, gray indicates additivity, and brown indicates antagonism) are shown for nineteen malignant cell lines grown as multi-cell type tumor spheroids and exposed to BI-3406 in combination with temuterkib. The malignant cell line name, tumor type, and KRAS status are indicated above each set of graphs.

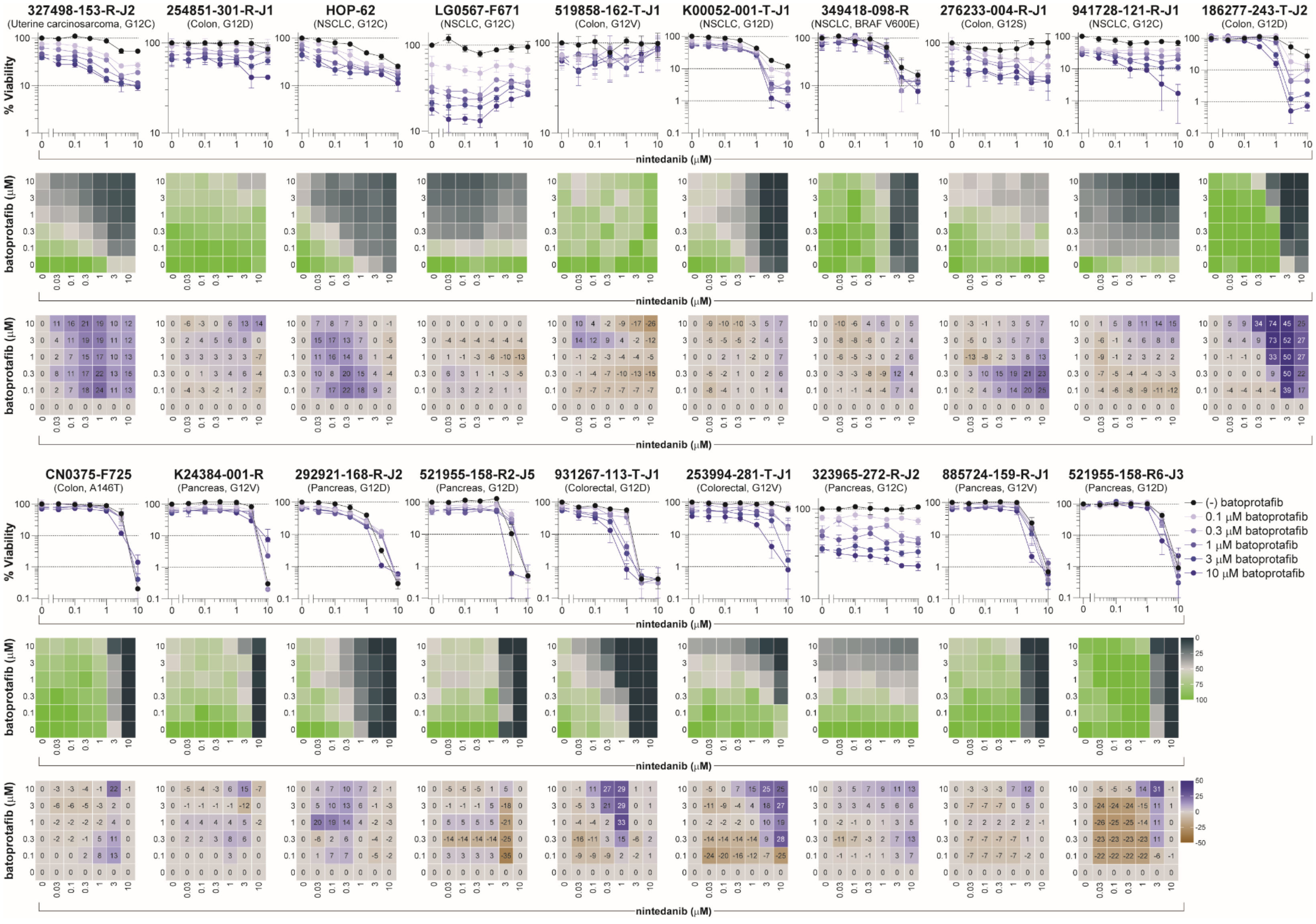


**Figure S6**. **Combination activity for batoprotafib with nintedanib in multi-cell type tumor spheroids**. Concentration-response graphs (*top*, mean ± SD, *n* = 4 technical replicates), % viability across the combination's concentration matrix (*middle*, mean of *n* = 4 technical replicates) displayed as a heatmap (green indicates high cell viability and black indicates low cell viability), and Bliss independence scores across the combination's concentration matrix (*bottom*, mean of *n* = 4 technical replicates) displayed numerically and as a heatmap (blue indicates synergy, gray indicates additivity, and brown indicates antagonism) are shown for nineteen malignant cell lines grown as multi-cell type tumor spheroids and exposed to batoprotafib in combination with nintedanib. The malignant cell line name, tumor type, and KRAS status are indicated above each set of graphs.

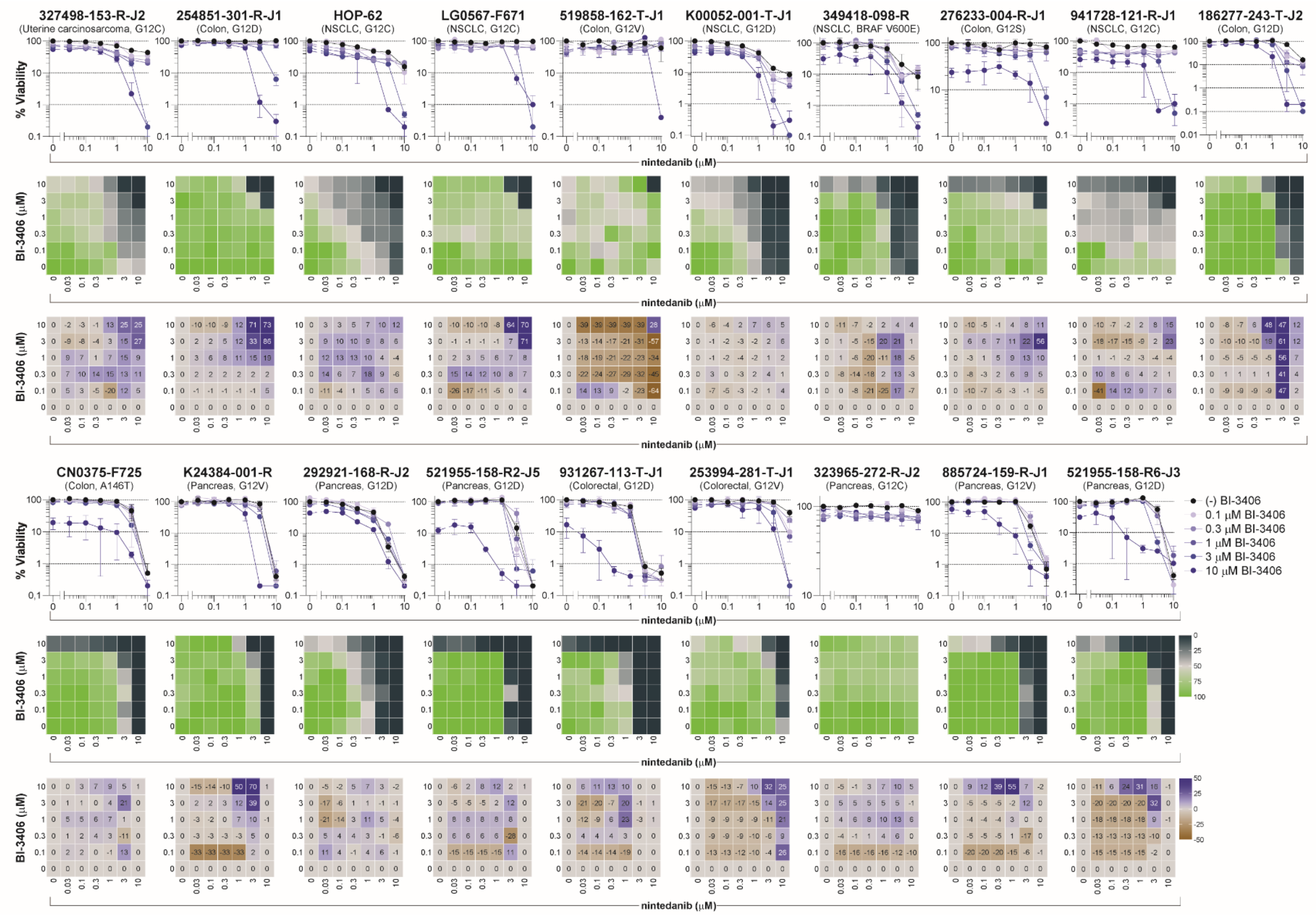

**Figure S7**. **Combination activity for BI-3406 with nintedanib in multi-cell type tumor spheroids**. Concentration-response graphs (*top*, mean ± SD, $n = 4$ technical replicates), % viability across the combination's concentration matrix (*middle*, mean of $n = 4$ technical replicates) displayed as a heatmap (green indicates high cell viability and black indicates low cell viability), and Bliss independence scores across the combination's concentration matrix (*bottom*, mean of $n = 4$ technical replicates) displayed numerically and as a heatmap (blue indicates synergy, gray indicates additivity, and brown indicates antagonism) are shown for nineteen malignant cell lines grown as multi-cell type tumor spheroids and exposed to BI-3406 in combination with nintedanib. The malignant cell line name, tumor type, and KRAS status are indicated above each set of graphs.

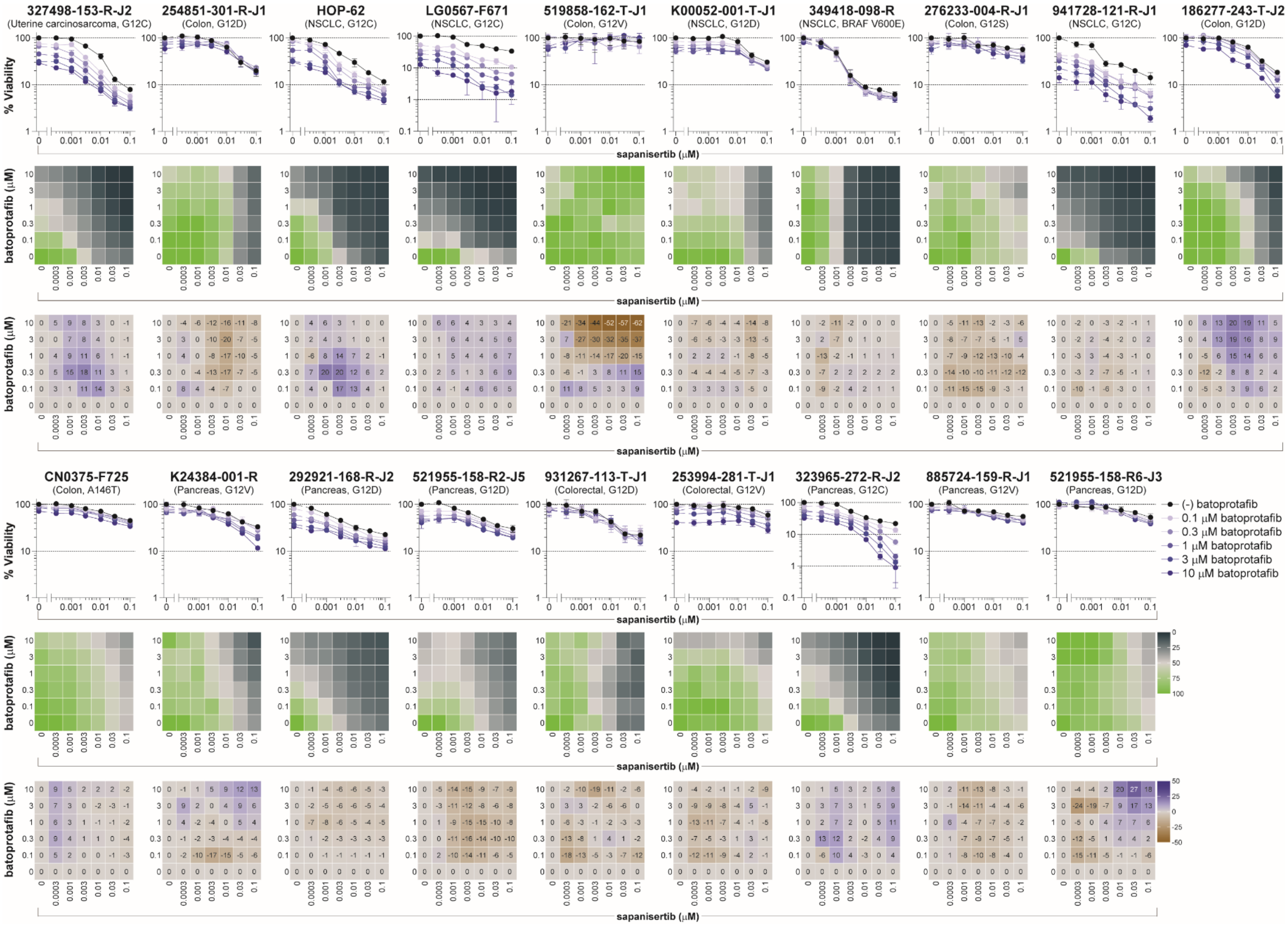


**Figure S8**. **Combination activity for batoprotafib with sapanisertib in multi-cell type tumor spheroids**. Concentration-response graphs (*top*, mean ± SD, *n* = 4 technical replicates), % viability across the combination's concentration matrix (*middle*, mean of *n* = 4 technical replicates) displayed as a heatmap (green indicates high cell viability and black indicates low cell viability), and Bliss independence scores across the combination's concentration matrix (*bottom*, mean of *n* = 4 technical replicates) displayed numerically and as a heatmap (blue indicates synergy, gray indicates additivity, and brown indicates antagonism) are shown for nineteen malignant cell lines grown as multi-cell type tumor spheroids and exposed to batoprotafib in combination with sapanisertib. The malignant cell line name, tumor type, and KRAS status are indicated above each set of graphs.

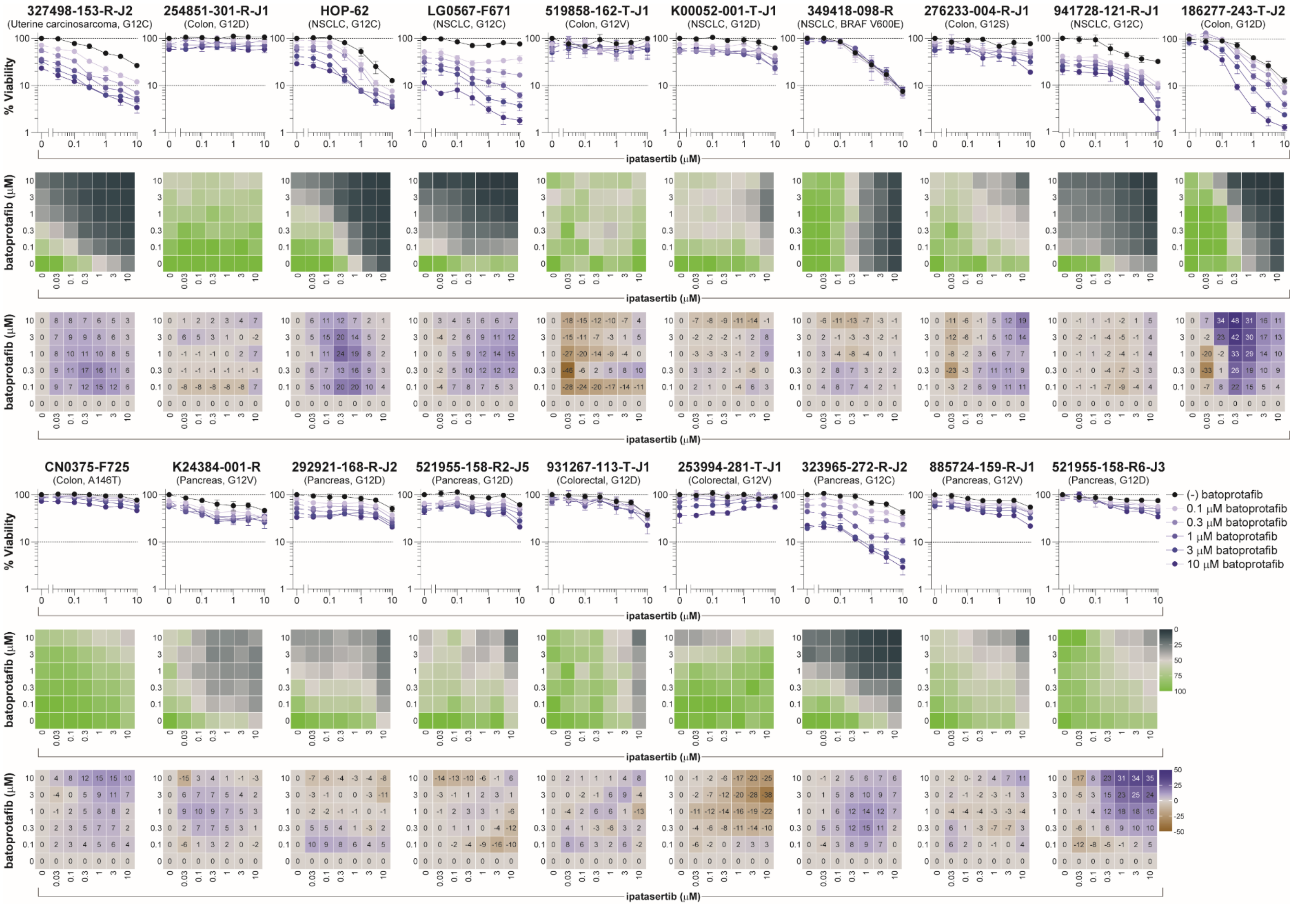


**Figure S9**. **Combination activity for batoprotafib with ipatasertib in multi-cell type tumor spheroids**. Concentration-response graphs (*top*, mean ± SD, *n* = 4 technical replicates), % viability across the combination's concentration matrix (*middle*, mean of *n* = 4 technical replicates) displayed as a heatmap (green indicates high cell viability and black indicates low cell viability), and Bliss independence scores across the combination's concentration matrix (*bottom*, mean of *n* = 4 technical replicates) displayed numerically and as a heatmap (blue indicates synergy, gray indicates additivity, and brown indicates antagonism) are shown for nineteen malignant cell lines grown as multi-cell type tumor spheroids and exposed to batoprotafib in combination with ipatasertib. The malignant cell line name, tumor type, and KRAS status are indicated above each set of graphs.

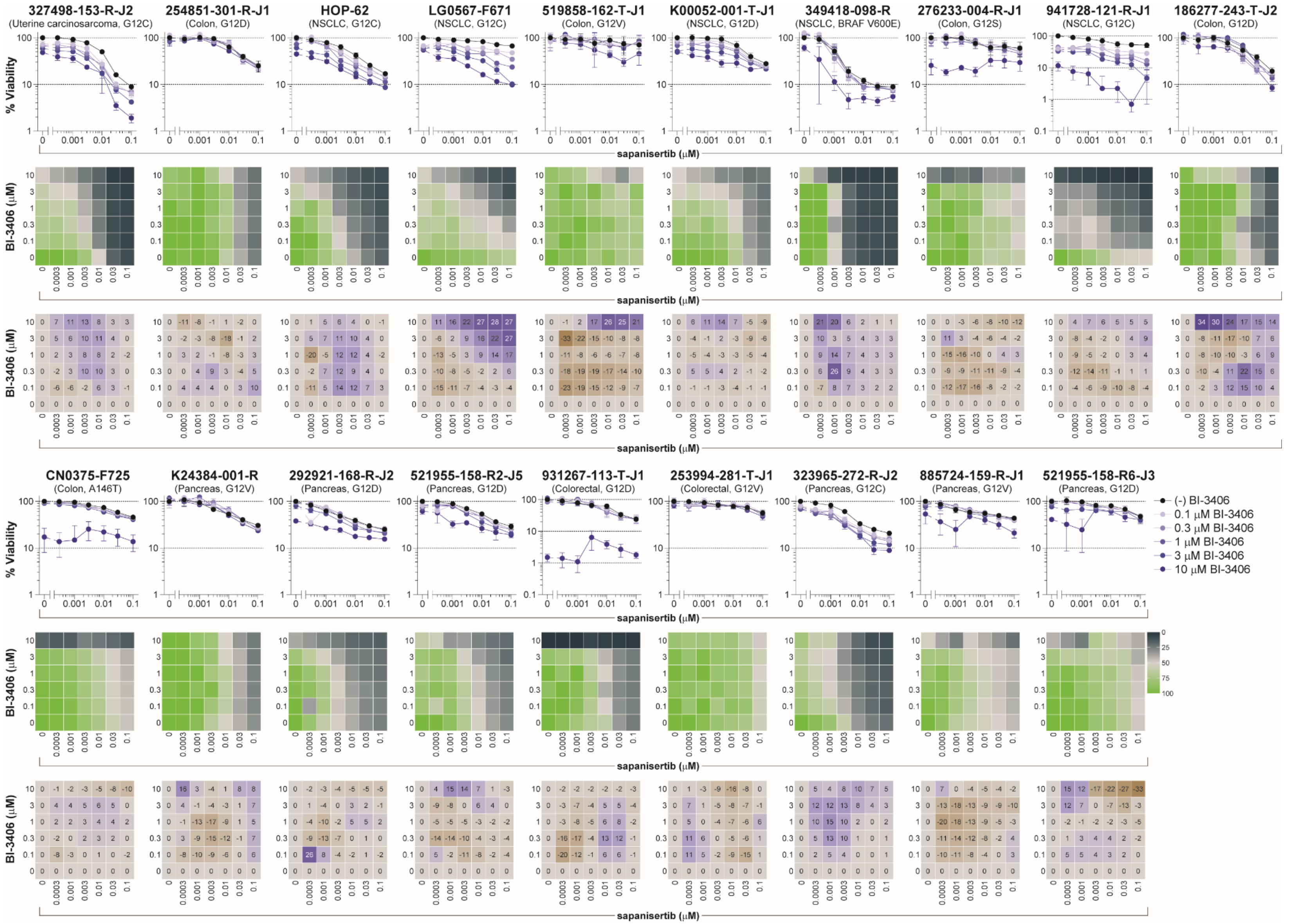


**Figure S10**. **Combination activity for BI-3406 with sapanisertib in multi-cell type tumor spheroids**. Concentration-response graphs (*top*, mean ± SD, $n = 4$ technical replicates), % viability across the combination's concentration matrix (*middle*, mean of $n = 4$ technical replicates) displayed as a heatmap (green indicates high cell viability and black indicates low cell viability), and Bliss independence scores across the combination's concentration matrix (*bottom*, mean of $n = 4$ technical replicates) displayed numerically and as a heatmap (blue indicates synergy, gray indicates additivity, and brown indicates antagonism) are shown for nineteen malignant cell lines grown as multi-cell type tumor spheroids and exposed to BI-3406 in combination with sapanisertib. The malignant cell line name, tumor type, and KRAS status are indicated above each set of graphs.

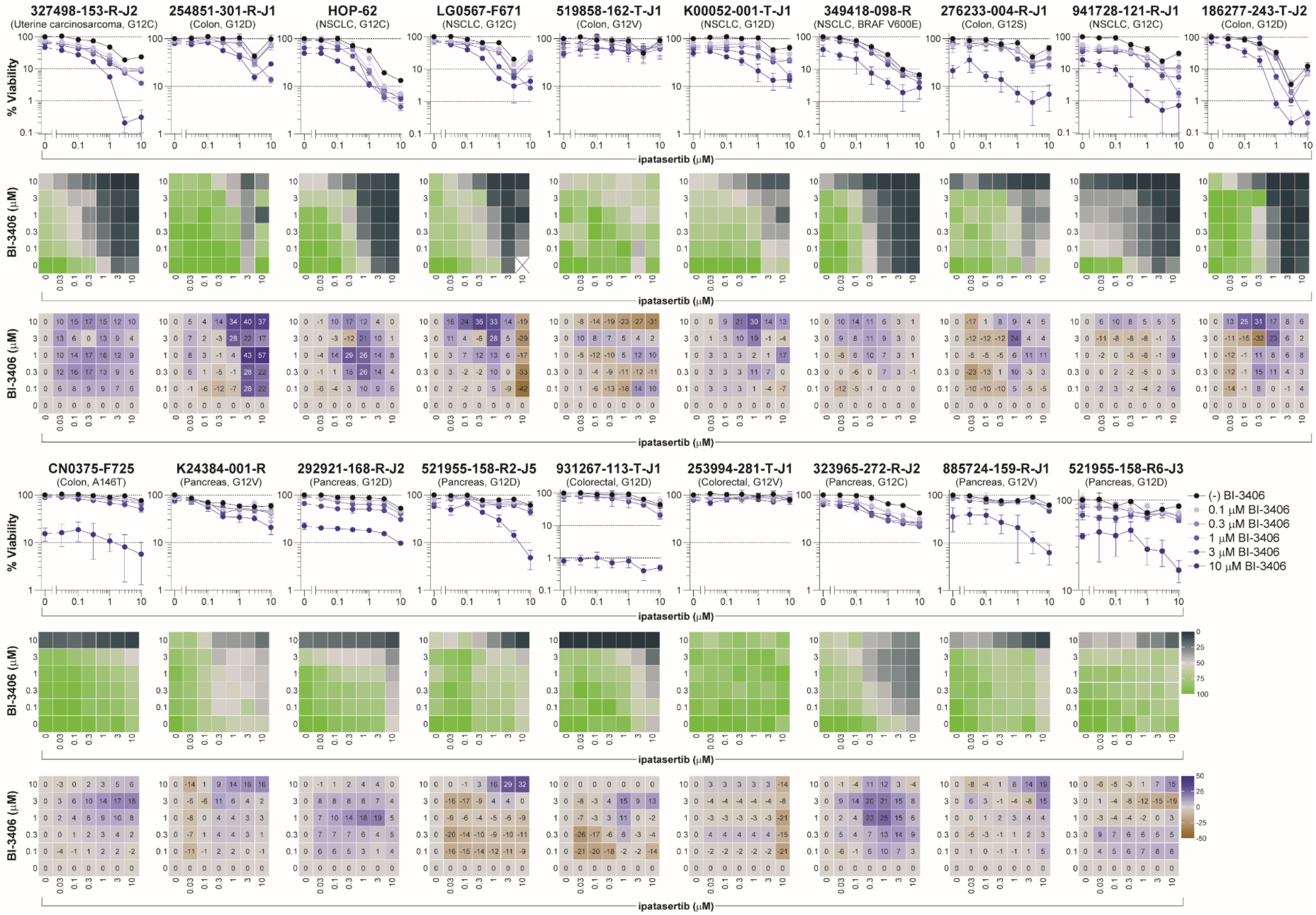


**Figure S11**. **Combination activity for BI-3406 with ipatasertib in multi-cell type tumor spheroids**. Concentration-response graphs (*top*, mean ± SD, $n$ = 4 technical replicates), % viability across the combination's concentration matrix (*middle*, mean of $n$ = 4 technical replicates) displayed as a heatmap (green indicates high cell viability and black indicates low cell viability), and Bliss independence scores across the combination's concentration matrix (*bottom*, mean of $n$ = 4 technical replicates) displayed numerically and as a heatmap (blue indicates synergy, gray indicates additivity, and brown indicates antagonism) are shown for nineteen malignant cell lines grown as multi-cell type tumor spheroids and exposed to BI-3406 in combination with ipatasertib. The malignant cell line name, tumor type, and KRAS status are indicated above each set of graphs.

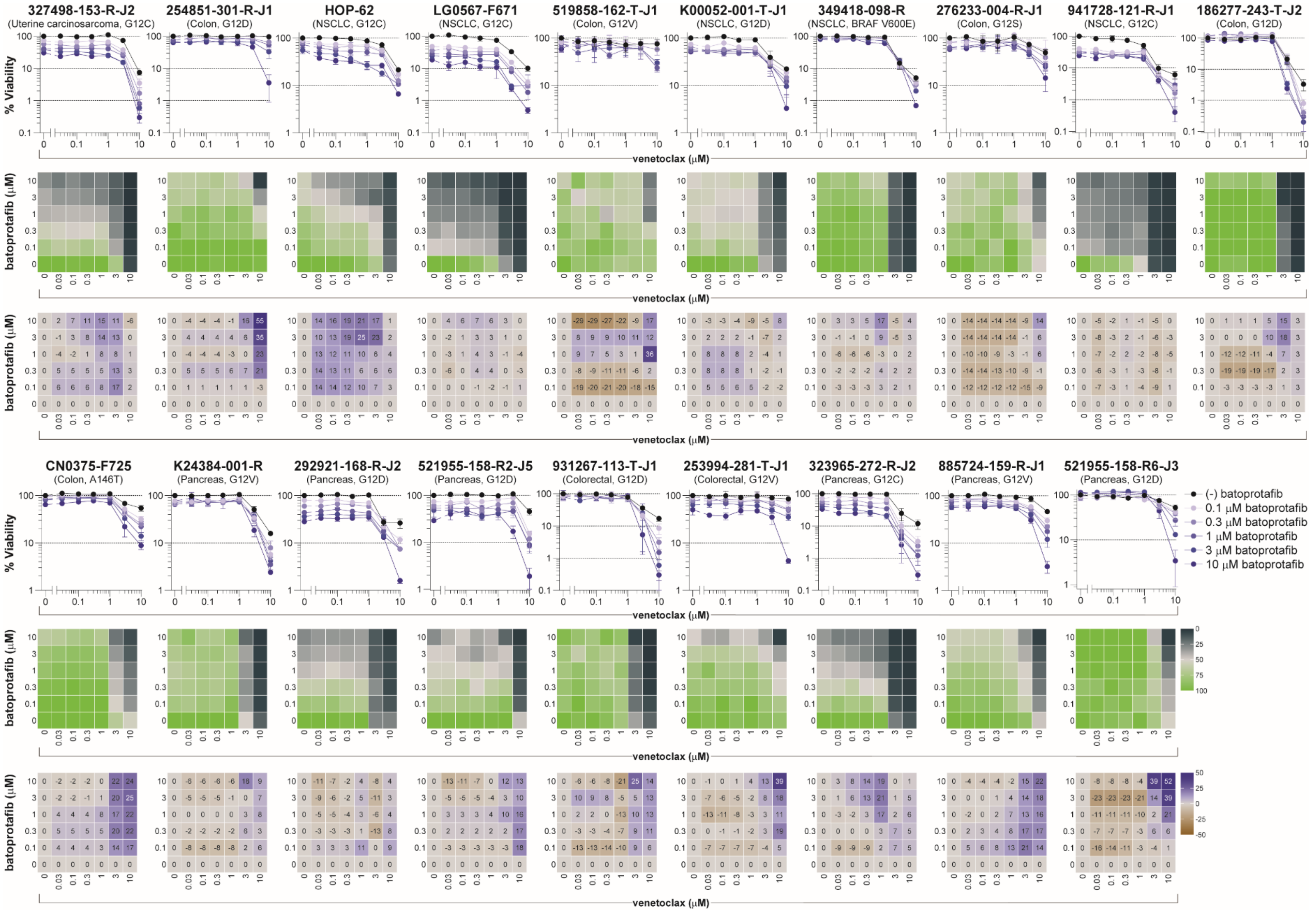


**Figure S12**. **Combination activity for batoprotafib with venetoclax in multi-cell type tumor spheroids**. Concentration-response graphs (*top*, mean ± SD, *n* = 4 technical replicates), % viability across the combination's concentration matrix (*middle*, mean of *n* = 4 technical replicates) displayed as a heatmap (green indicates high cell viability and black indicates low cell viability), and Bliss independence scores across the combination's concentration matrix (*bottom*, mean of *n* = 4 technical replicates) displayed numerically and as a heatmap (blue indicates synergy, gray indicates additivity, and brown indicates antagonism) are shown for nineteen malignant cell lines grown as multi-cell type tumor spheroids and exposed to batoprotafib in combination with venetoclax. The malignant cell line name, tumor type, and KRAS status are indicated above each set of graphs.

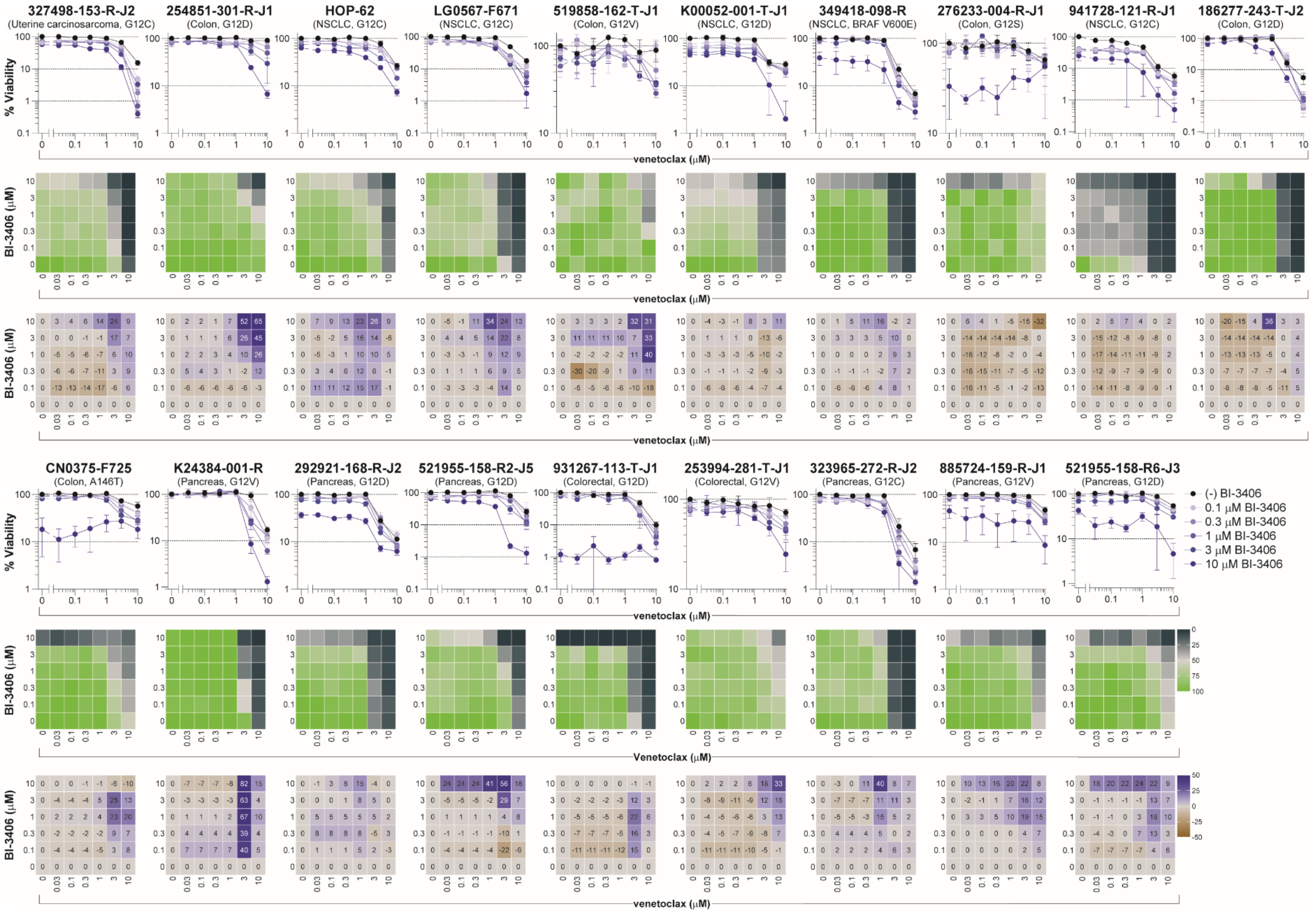


**Figure S13**. **Combination activity for BI-3406 with venetoclax in multi-cell type tumor spheroids**. Concentration-response graphs (*top*, mean ± SD, *n* = 4 technical replicates), % viability across the combination's concentration matrix (*middle*, mean of *n* = 4 technical replicates) displayed as a heatmap (green indicates high cell viability and black indicates low cell viability), and Bliss independence scores across the combination's concentration matrix (*bottom*, mean of *n* = 4 technical replicates) displayed numerically and as a heatmap (blue indicates synergy, gray indicates additivity, and brown indicates antagonism) are shown for nineteen malignant cell lines grown as multi-cell type tumor spheroids and exposed to BI-3406 in combination with venetoclax. The malignant cell line name, tumor type, and KRAS status are indicated above each set of graphs.
